# Connectome-based spatial statistics enabling large-scale population analyses of human connectome across cohorts

**DOI:** 10.64898/2026.04.09.717492

**Authors:** Tengfei Li, Xifeng Wang, Martin Cole, Zehui Sun, Zhiwen Jiang, Xinjie Qian, Shan Gao, Tianyou Luo, Maxime Descoteaux, Jason L. Stein, Xiao Wang, Thomas E. Nichols, Heping Zhang, Zhengwu Zhang, Hongtu Zhu

## Abstract

Large-scale population analyses of structural connectome organization remain challenging because of cross-subject alignment, pathway interpretability and computational burden. No widely adopted standard exists for systematic evaluation across processing methods. We developed connectome-based spatial statistics (CBSS), a scalable framework for anatomically aligned and functionally informed quantification of white-matter microstructure that yields atlas-defined pathways organized into *13* functional networks. Using data from *56,510* UK Biobank participants together with five independent lifespan cohorts, we evaluated the streamline-, voxel– and network-level measures in the aspects of reliability, heritability, structure–function coupling, cognitive and behavioral prediction, brain aging patterns and lifespan trajectories across cohorts. The systematic evaluation workflow compares population-level white-matter representations across methods, spatial scales, tasks and datasets. The results support CBSS as a common connectome reference for large-scale, cross-cohort diffusion MRI studies.

## Main

Diffusion MRI (dMRI) has become a cornerstone for in vivo mapping of human white matter (WM) and structural connectivity. Among the current dMRI analysis methods, tract-based spatial statistics (TBSS) ^1^ has been widely applied to large dMRI datasets for the two decades, because it is fast, robust, spatially aligned, and compatible with various acquisitions with a user-friendly tract– and voxel-based framework. To better accommodate multiple fiber orientations within a voxel, fixel-based spatial statistics (FBSS) ^2^ further introduced population inference on fixel-wise measures. However, those voxelwise frameworks are not explicitly organized around the WM connectome and therefore provide limited pathway-level interpretability and network-level context.

By contrast, tractography-based approaches ^3–5^ reconstruct whole-brain streamline architectures, enabling pathway-level quantification, connectome construction and network analyses. A range of tractography atlases and atlas-guided frameworks ^6–12^ have been proposed, including fiber clustering-based atlases, population-averaged tract-to-region connectome, tractography atlas-based spatial statistics, functional MRI-guided probabilistic WM atlases, and atlas prior-constrained pathway reconstruction. However, large-scale population analyses remain challenging because whole-brain tractography and downstream pathway delineation can be computationally intensive, and estimated pathway geometry and correspondence can vary with acquisition, preprocessing and tractography choices. These factors can limit throughput and cross-cohort reproducibility at the biobank scale.

To this end, we introduce Connectome-Based Spatial Statistics (CBSS), a framework that leverages a brain function–aware structural connectome (BFSC) atlas as a common population reference for rapid mapping and analysis of diffusion measures. Harnessing massive-scale tractography encompassing billions of streamlines from *1,042* high-quality Human Connectome Project Young Adult (HCP-Y) ^13^ participants, we constructed BFSC by integrating cortical-surface registration, function-aware connectome parcellation, and fiber clustering, yielding fine-grained atlas pathways with explicit functional-network context. CBSS allows mapping diffusion measures from heterogeneous cohorts to BFSC atlas using fine-grained projection and along-path alignment to improve robustness, and enables voxel-, pathway– and network-level traits within a unified framework. By avoiding subject-level whole-brain tractography in downstream cohorts, CBSS largely reduces computational burden and facilitates biobank-level diffusion data analyses.

To demonstrate the robustness, scalability, and biological validity of CBSS, we systematically evaluated the framework across more than *64,000* diffusion scans from the UK Biobank and five independent lifespan cohorts (**Figure 1**). Because a widely adopted standard for evaluating dMRI processing and connectome quantification is currently lacking, we assembled a unified evaluation workflow spanning a broad array of analysis contexts. Using this comprehensive approach, we establish that CBSS provides exceptional test-retest reliability, captures strong heritability and structure–function coupling, and robustly predicts cognitive and behavioral traits. Moreover, our multi-cohort lifespan design enables rigorous cross-cohort validation, offering an integrated picture of variations in white-matter integrity and connectome organization across all phases of human life. Ultimately, these results validate CBSS as a powerful, functionally interpretable, and computationally efficient common reference framework for large-scale, cross-cohort structural connectomics.

**Figure 1.**
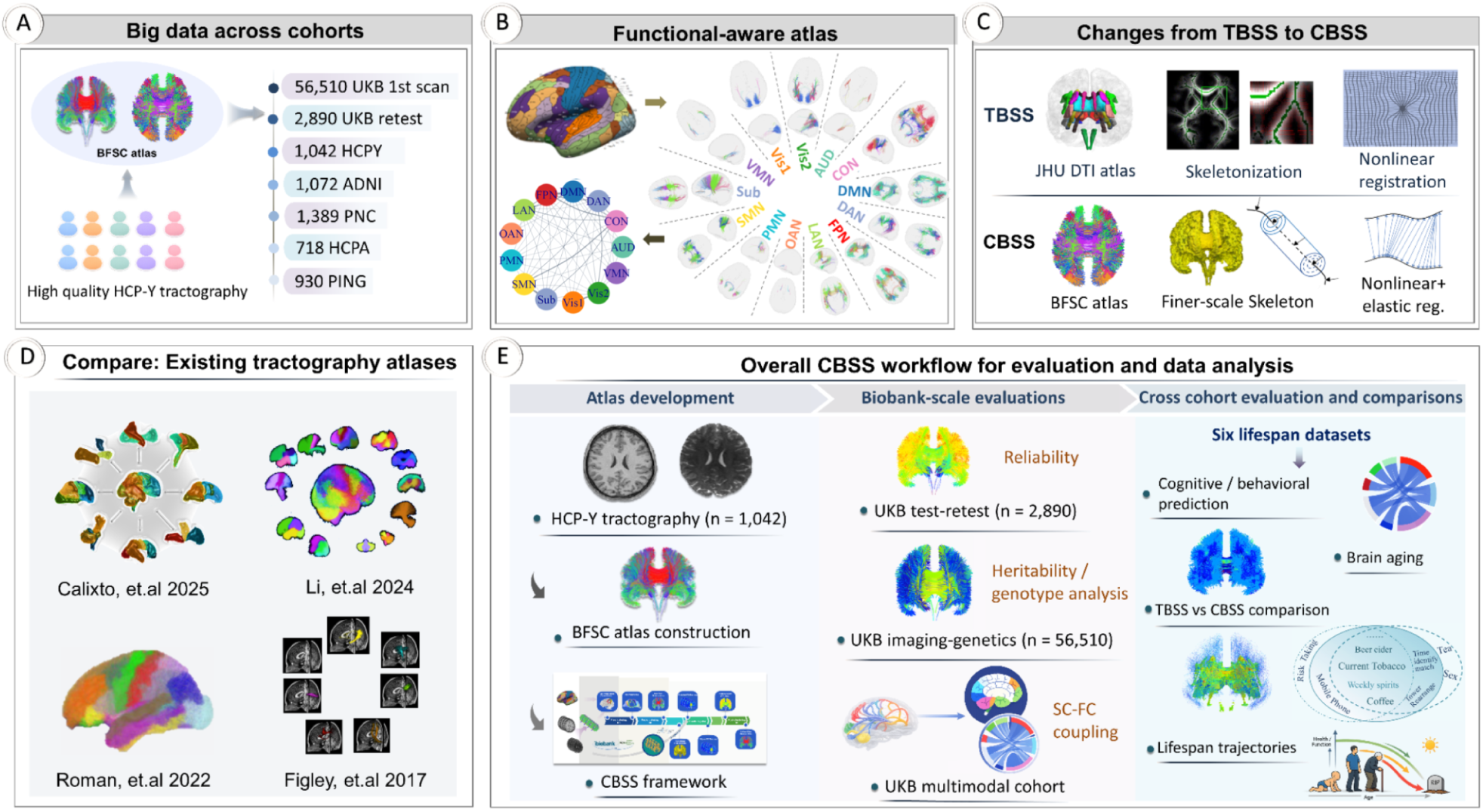
The study design. **A.** The use of six multi-site datasets in the study (7 datasets, 64,551 diffusion scans). **B.** Network visualization of the BFSC atlas and connectome. Cortical parcellation into 13 functional networks (upper left) defines fiber endpoints; the right panel shows WM pathways linking from a given functional network parcel to all the others. These pathways can quantify between network structural connectivity based on fiber counts (lower left). **C.** Major differences between the CBSS and TBSS pipelines: atlas, diffusion projection, and registration methods. **D.** Other existing tractography atlases ^8,9,11,12^. **E.** Overall CBSS workflow for evaluation and data analysis. BFSC, brain function–aware structural connectome (atlas); TBSS, tract-based spatial statistics; CBSS, connectome-based spatial statistics; SC, structural connectivity; FC, functional connectivity. 13 networks, including AUD (auditory), DMN (default mode), FPN (frontoparietal), DAN (dorsal attention), CON (cingulo–opercular), SMN (somatomotor), LAN (language), OAN, (orbito-affective), VMN (ventral multimodal), PMN (posterior multimodal), Vis1 and Vis2 (primary and secondary visual), and Sub (subcortical) networks.

## Results

### Overview of atlas construction and CBSS mapping

We constructed the BFSC atlas leveraging high-quality dMRI and structural MRI (sMRI) from *1,042* HCP-Y participants (**Figure 1A**; **Online Methods**; **Supplementary Figure S1**). Whole-brain tractography was generated ^3^ for each subject and anchored to a fine-grained *359*-region parcellation. This parcellation was derived from the HCP-MMP cortical atlas ^14^ and *13* FreeSurfer subcortical regions, with selected adjacent cortical parcels merged to improve tractography clustering stability (**Supplementary Figure S2**). These regions were subsequently organized ^15^ into 13 functional networks encompassing sensory, cognitive-control, multimodal and affective systems (**Figure 1B)**. Using HCP test–retest data, we evaluated fiber-count reliability between region pairs to ensure only highly reproducible pathways were retained for atlas construction. Subject-level tractography was then aligned to the MNI152 reference space by non-linear alignment and aggregated across subjects into a population-scale tractogram. Functional-network–aware clustering with TractDL ^16^ yielded *6,090* pathway clusters, of which *5,723* cluster centroids were retained after quality control to define the final BFSC atlas.

With the BFSC atlas established as a normative reference, the CBSS framework can seamlessly map diffusion-derived measures from new cohorts to this atlas without requiring computationally heavy subject-level whole-brain tractography. To achieve this, subject diffusion maps are nonlinearly registered to MNI152 space, projected onto representative pathways using a fine-scale skeletonization and projection procedure ^1^, and further refined via elastic registration ^17^ to reduce along-path shifts (**Figure 1C**).

### The BFSC atlas

The atlas retained *5,723* representative WM pathways, categorized into *2,231* within-network and *3,492* between-network connections that span *68* functional-network pairs ^15^ (**Figures 1B**, **2A** and **Supplementary Figure S3**). Evaluation of the network-level pathway distribution reveals substantial heterogeneity (**Supplementary Figure S4**). For instance, the default mode (DMN) and frontoparietal (FPN) networks contribute the highest number of pathways—including densely represented DMN–DMN and DMN–FPN connections—whereas others, such as the ventral multimodal (VMN) and primary visual (Vis1) networks, lack within-network pathways following stringent quality control.

Geometric profiling of the atlas demonstrates a diverse representation of pathway shapes and sizes (**Supplementary Figure S5**). **Figure 1B** and **2A (**left) visualize representative pathways across network pairs. Mean pathway length and curvature are strongly inversely correlated (*r = – 0.81;* **Figure 2A**, **right**); shorter, more curved pathways typically connect anatomically adjacent networks, while longer, straighter pathways link spatially distributed network pairs.

**Figure 2.**
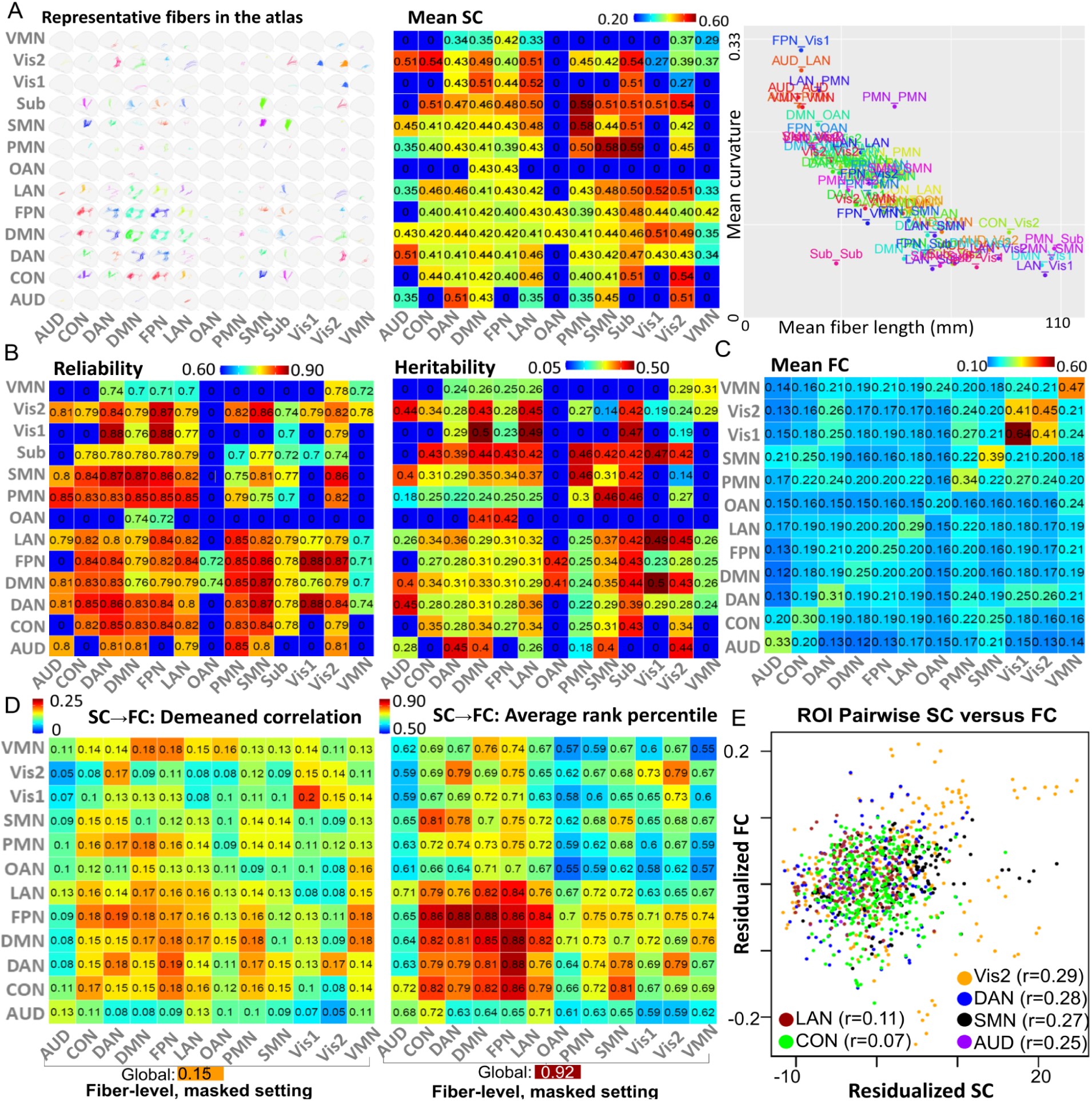
Common network attributes of the BFRC atlas across 13*13 network pairs. **A.** Spatial location (left), population-mean FA-based structural connectivity (middle), and mean fiber length (right: x-axis) versus mean curvature (right: y-axis). **B.** The mean pathway-level test-retest reliability (left) and heritability (right). In A and B, *0* indicates absence of a pathway in the atlas. **C.** The functional-network level averaged functional connectivity. **D.** Network pairwise summaries of FC prediction performance using pathway-average FA: average demeaned correlation (left) and rank percentile (right). **E.** Population-mean FC versus SC for six networks with the strongest SC-FC coupling (correlation *r* across ROI pairs involving each network). ROI pairwise SC and FC were residualized by regressing out log-transformed fiber length (**Supplementary Figure S9B**), and each point represents the (SC, FC) between one ROI pair from the HCP-MMP atlas. ROI pairs connecting two networks were plotted twice in two networks colors, with small offsets added for visualization. SC/FC, structural/functional connectivity; FA, fractional anisotropy; ROI, regions of interest.

Finally, to assess anatomical validity, we measured overlap between BFSC pathways and the classical JHU ICBM-DTI-81 atlas (**Supplementary Figure S6**). Corpus callosum overlap was segment-specific, with the genu dominated by DMN-centred interhemispheric pathways, the body by language-, cingulo-opercular-, somatomotor– and frontoparietal-related pathways, and the splenium by posterior multimodal (PMN)-related pathways. The posterior thalamic radiations were associated mainly with visual-centred connections, the posterior limb of the internal capsule with subcortical–cortical pathways, and the superior longitudinal fasciculus with association pathways linking the PMN, somatomotor (SMN), dorsal attention (DAN) and FPN systems.

### Connectivity, test-retest reliability, and heritability

To evaluate the reproducibility and genetic basis of the BFSC atlas, we applied the CBSS framework at scale using data from UK Biobank ^18^. Specifically, we leveraged 2,890 participants with baseline and repeat dMRI to estimate test–retest reliability (**Figure 2B, left**), and 56,510 participants with both baseline dMRI and genome-wide single-nucleotide polymorphism (SNP) data to quantify heritability **(Figure 2B, middle** and **Supplementary Figure S7**). Baseline structural connectivity (SC) among the 13 functional networks, defined by fractional anisotropy (FA)-weighted edge strength, is highest for connections involving the subcortical (Sub), posterior multimodal (PMN), and SMN networks. We also observed notably high connectivity for pathways linking the secondary visual (Vis2) and auditory (AUD) systems with higher-order association and control networks (**Figure 2A**, **middle**).

Across network-pair edges, test–retest reliability is consistently high (mean intraclass correlation coefficient [ICC] *= 0.81 ± 0.056,* **Figure 2B, left**), with most edges falling in the range of *0.75-0.88*. Relatively lower reliability is confined to a subset of edges involving subcortical and ventral multimodal systems (ICC *= 0.70-0.74*), whereas edges involving higher-order association and control networks show broadly strong reproducibility. These spatial patterns are consistent with prior reports ^3,19,20^ that diffusion-derived microstructural measures are generally highly reliable, with tract– and network-dependent variation.

Furthermore, the SNP-based heritability of these FA-weighted network edges is moderate to high across the majority of the network matrix (*h*^2^ *0.18-0.50,* **Figure 2B, middle**). The highest heritability estimates are observed for several subcortical–cortical and association–sensory couplings. In particular, edges linking the DMN, language (LAN), and FPN networks with subcortical and visual systems emerge as the most heritable, alongside selected auditory–association connections (*h*^2^ *0.40–0.45*). These magnitudes and spatial patterns are highly concordant with previous large-scale dMRI ^21,22^ and structural connectome studies ^23–25^, reinforcing the substantial contribution of common genetic variants to white-matter microstructural organization.

### Structural-functional coupling

Having established that CBSS captures reliable and heritable structural features, we next investigated the extent to which this anatomical connectome aligns with brain function. For each subject, we derived a *360* × *360* functional connectivity (FC) matrix using the UK Biobank preprocessing pipeline ^26^ and the HCP-MMP cortical parcellation. We retained *64,620* unique region-to-region FC edges for structural-to-functional connectivity (SC-FC) prediction (**Figure 2C**). Using partial least squares (PLS), we predicted these FC edges from CBSS-derived SC features at the fiber, voxel, and network levels. To ensure robust generalization, models were trained on *19,199* UK Biobank Phase 1–2 participants, tuned on *3,088* Phase 4 participants, and evaluated in an independent Phase 3 test set of *18,136* participants (**Figure 3A**).

**Figure 3.**
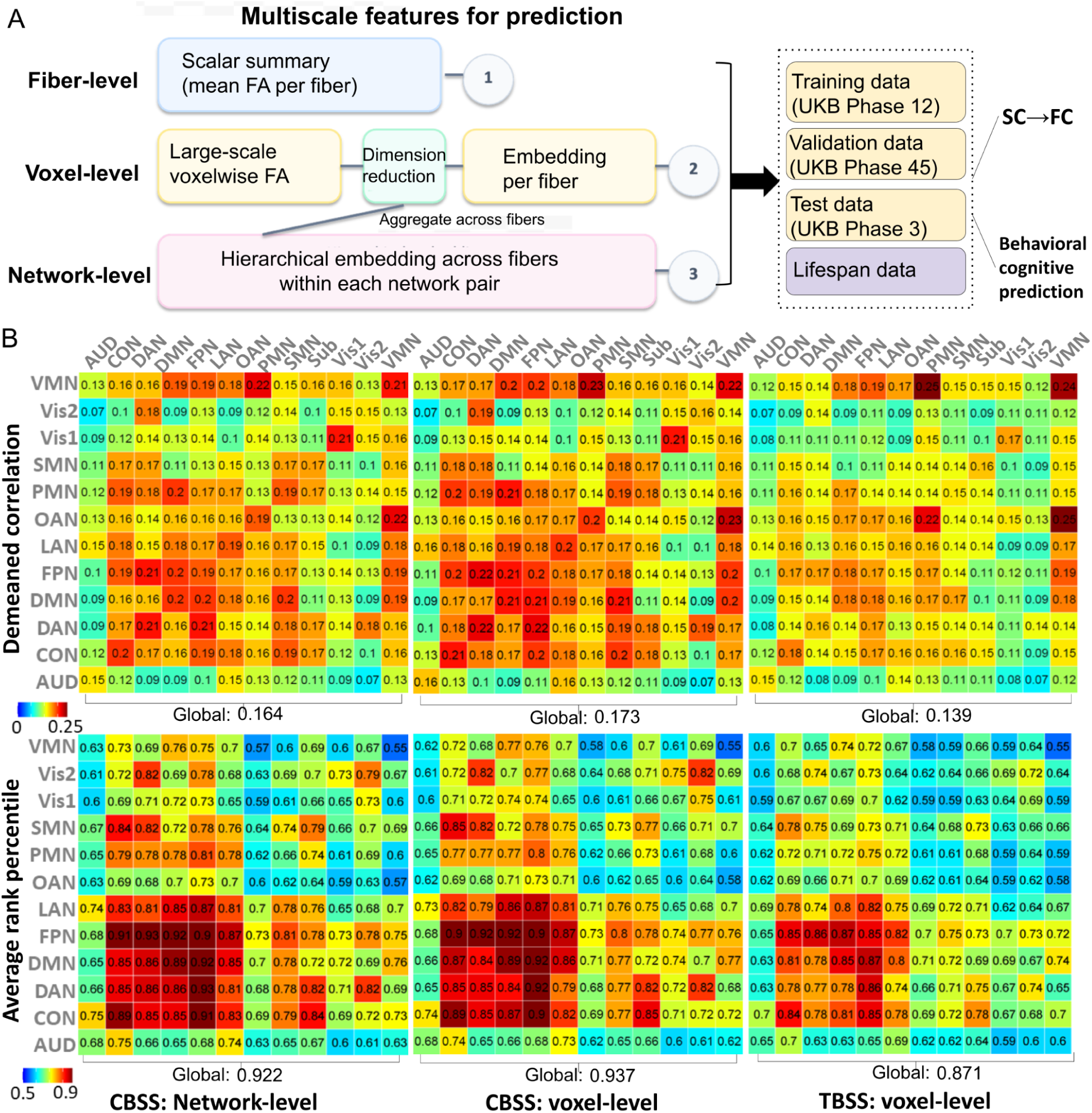
Multiscale features for prediction tasks. **A.** Schematic overview for tasks of predicting functional connectivity, and cognitive and behavioral phenotypes. CBSS feature constructions based on fiber-, voxel-, and network-level embeddings (principal components) and summary statistics via the PLS prediction model and the evaluation design of UK Biobank and external lifespan datasets. **B.** Brain structural functional coupling results, where de-meaned correlation and average rank percentile for FC predictions were compared among the TBSS and the network-level and voxel-level CBSS across all network pairs. SC, structural connectivity; FC, functional connectivity; PLS, partial least square regression.

For fiber-level SC feature construction, we evaluated two conditions: a non-masked setting using all atlas pathways as predictors, and a masked setting restricted to pathways in which at least one endpoint shared the same functional network with the target FC edge. Performance was summarized at both the network-pair level and global level using the de-meaned correlation (*avgcorr*) and average rank percentile (*avgrank*), following *Jamison, et al*. ^27^. Specifically, at the network-pair level, for each subject, we computed the de-meaned Pearson correlation between predicted and observed FC patterns across pairs of regions of interest (ROIs) in HCP-MMP parcellation within that network pair, and then averaged this quantity across subjects. At the whole-brain level, we computed the same correlation across all *64,620* ROI pairs and again averaged across subjects. Similarly, *avgrank* was defined as the percentile rank of each subject’s predicted–observed FC correspondence relative to the corresponding cross-subject matches. This was computed using FCs within each network pair at the network-pair level, or using all ROI-pair FCs at the whole-brain level, and then averaged across subjects. In the non-masked setting, SC-FC coupling was widespread but system-dependent (global *avgcorr* = *0.15* and *avgrank* = *0.90*), with peak performance in association, control, and attention systems, including LAN, FPN, DMN, DAN, and orbito-affective (CON) networks (**Supplementary Figure S8**). In the masked setting, restricting predictors to network-relevant pathways improved identifiability (global *avgrank* = *0.92*; **Figure 2D**).

To quantify SC contributions to FC prediction, we defined a hierarchical contribution measure: for each fiber pathway, its contribution to a specific ROI-pair FC was defined as the cross-subject correlation between pathway-averaged FA and predicted FC value for that ROI pair; the contribution of that pathway to an FC network pair was then defined as the maximum of these correlations across all ROI pairs within the network pair. We used the maximum as a better-match summary than mean, because a given structural pathway is expected to align most strongly with only a subset of ROI-pair FCs within a network pair, and the maximum can pick the best match. SC contributions were strongly network-concordant in the auditory, visual, and attention systems, where FC was predicted primarily by SC from the corresponding networks, indicating high SC–FC coupling in these domains (**Supplementary Figure S9A**). Consistent with the prediction results, ROI-pairwise SC–FC correlations were strongest in sensory networks (Vis2, SMN and AUD) and in the dorsal attention network (DAN), relative to other higher-order cognitive networks (**Figure 2E**). Because the contour analysis (**Supplementary Figure S9B**) revealed a nonlinear dependence of population-mean FC on SC and fiber length, the SC and FC values in **Figure 2E** were residualized for log-transformed fiber length.

We next compared SC–FC prediction in the UK Biobank phase-3 test set using CBSS voxel– and network-level embeddings and TBSS voxel-level embeddings, all under the same non-masked setting as above for a matched comparison. TBSS achieved *avgcorr* = *0.139* and *avgrank* = 0.*871*, whereas CBSS increased *avgcorr* to *0.164* at the network-level and *0.173* at the voxel-level, with corresponding *avgrank* of *0.922* and *0.937* (**Figure 3B**). Compared with both fiber-level CBSS and voxel-level TBSS, voxel– and network-level CBSS improved spatial correspondence and subject identifiability, indicating that voxel-level and hierarchical network-level embeddings capture additional subject-specific information beyond tract averaging. The largest gains were observed in higher-order association and control systems, including FPN, CON, DAN and DMN. Under the same evaluation metrics, these values exceeded the SC→FC benchmark reported by *Jamison, et al.* ^27^ (*avgcorr* = *0.12*, *avgrank* = *0.91*), although it used fiber-count-based SC.

### Cognitive and behavioral predictions

We predicted *96* UK Biobank cognitive and behavioural phenotypes (**Supplementary Table 1**) using connectome features defined at the CBSS-fiber, voxel and network levels, following the procedure in **Figure 3A**. In the UKB phase-3 hold-out test set, PLS models based on voxel-level features outperformed those based on fiber-level summaries for most phenotypes (**Supplementary Figure S10**), indicating that intra-tract spatial variation carries predictive signals beyond tract-averaged measures. The largest improvements were observed for the strongest targets: sex predict accuracy improved from *0.928* to *0.951* and age correlation from *r* = *0.758* to *0.786*. Prediction of a leading cognitive phenotype, digit matching, also improved from *0.357* to *0.388*, with smaller but directionally consistent gains across many additional traits.

We next performed network-specific prediction, by fitting each of the above prediction models to fiber-level FA features from pathways with at least one endpoint in a given network. Predictive signal was shown to be broadly distributed across connectome (**Figure 5A**, left). Cognitive and education-related phenotypes were most strongly represented in visual and frontoparietal, attention and control systems, whereas demographic and lifestyle phenotypes involved a broader anatomical substrate, including DMN and subcortical systems. Because the final PLS predictor is a linear combination of FA features with both positive and negative regression coefficients, we decomposed it into positive– and negative-coefficient contributions to aid interpretation ^28^. Specifically, after model fitting, we separated the final regression coefficients by sign and projected the corresponding FA features onto each subset to obtain signed prediction components (**Supplementary Figure S11**). This decomposition provides a descriptive summary of which signed part of the fitted multivariate predictor contributed more strongly to each phenotype. Age, smoking, alcohol intake, television viewing, oily fish intake and lamb/mutton consumption were driven primarily by the negative-coefficient components, whereas education, fluid intelligence, phone use, driving and miserableness were driven primarily by the positive-coefficient component. Other phenotypes, including sex, race, bread intake, processed meat intake, neuroticism score and risk-taking, showed mixed contributions. Directions of the top contributors for each network (**Supplementary Figure S12**) further summarized, emphasizing this signed organization. Together, these results suggest that individual differences in cognition and behaviour are encoded not simply by the magnitude of structural connectivity, but by a distributed and signed reorganization of network-level connectivity across the connectome.

We compared between CBSS and TBSS using matched feature extractions and modelling pipelines (**Figure 3A**) for cognitive and behavioral trait predictions in UKB and other five lifespan cohorts (**Figures 4A-C**, and **Supplementary Figure S13**). CBSS generally exceeded TBSS on the UKB phase 3 hold-out set in prediction of race, skin color, and hair colors, while the “CBSS+TBSS” model utilizing the combined features could achieve the best predictive performance across the majority of phenotypes (**Figures 4B and 4C**). External validation across lifespan cohorts showed CBSS significantly improves prediction for cognition-related phenotypes. We also implemented and compared two predictive models: the PLS and elastic net (EN). In HCP-Y, CBSS improves early, fluid and total intelligence scores from *0.14*, *0.19* and *0.25* to *0.30*, *0.29*, and *0.34*. In HCPA, CBSS improves early, crystal and total intelligence scores from *0.50*, *0.25* and *0.43* to *0.58*, *0.41*, and *0.53*. Similar CBSS gains extend to clinical cognition in ADNI, such as ADAS13 from *0.48* to *0.52*. With the exception of ADNI ADAS-score prediction and the TBSS method with HCP-A data, PLS generally outperformed elastic net (EN). Performances to validate UKB model transfer are shown in **Supplementary Figure S13**. Overall, these results indicate that CBSS, especially its network/voxel-level features, provides more comprehensive white matter information in cognitive and behavioral prediction tasks.

**Figure 4.**
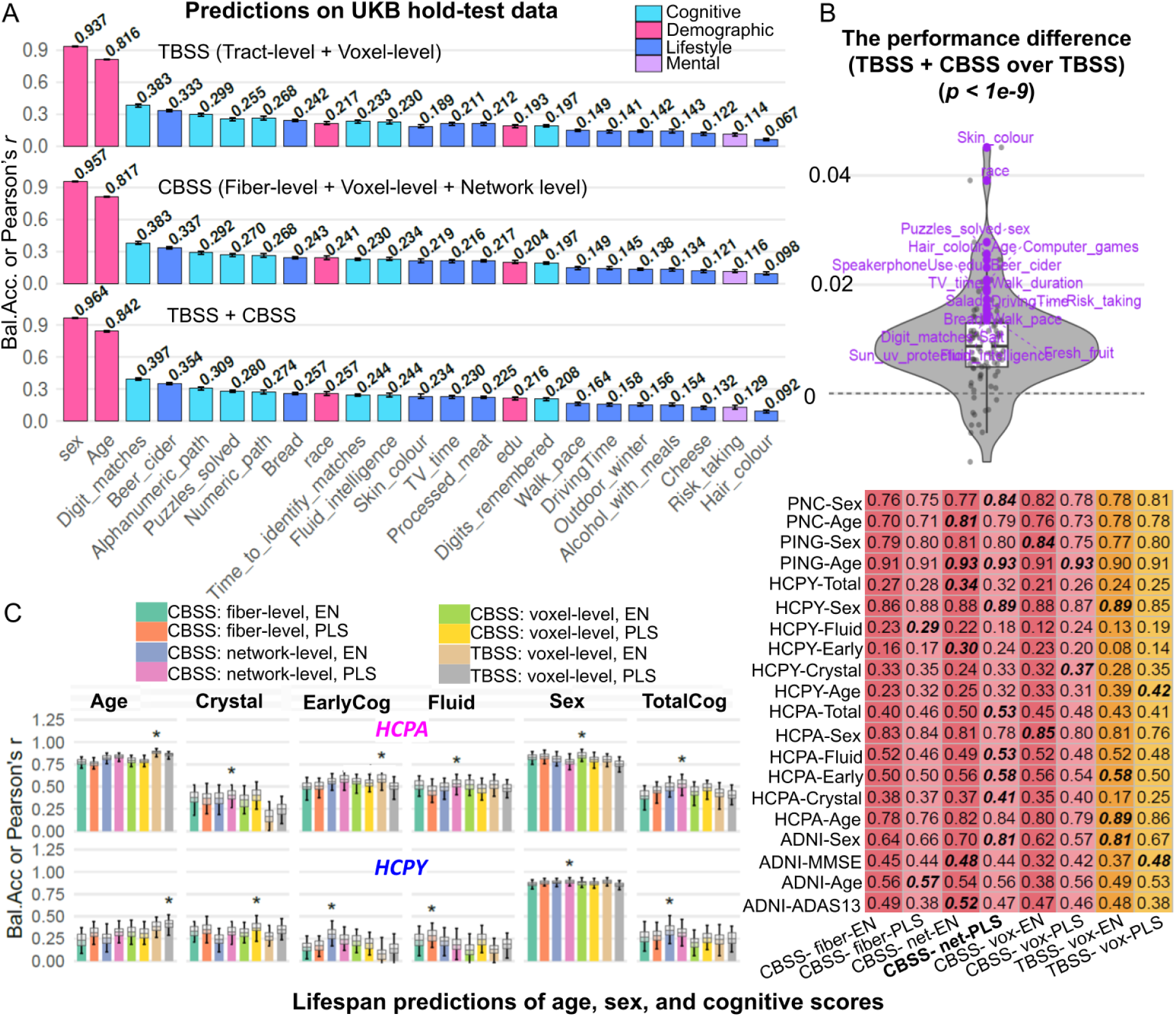
Prediction comparisons of age, sex, and cognitive/behavioral performances. **A.** UK Biobank hold-out performance (balanced accuracy for sex; Pearson’s *r* for continuous traits) using TBSS, CBSS, and TBSS+CBSS feature sets (TBSS: tract+voxel; CBSS: fiber/pathway+voxel+network), based on an ensemble of EN and PLS models. Colors indicate trait categories and values are annotated above bars. **B.** Per-trait performance difference of TBSS+CBSS over TBSS on the UK Biobank hold-out set. P values were computed using a paired t-test to assess the global significant difference across all traits. **C**. Hold-out performance on five lifespan datasets using CBSS fiber-, voxel– and network-level features and TBSS voxel-level features, evaluated with EN and PLS, separately. Bar plots (left) show mean performance with 95% confidence intervals and asterisks mark the best-performing approach; the right table summarizes mean performance across cohorts. The top performing approach in each row is shown in bold texts. *Bal.Acc*, balanced accuracy; Crystal, crystallized intelligence; Fluid, fluid intelligence; TotalCog, total cognition; EarlyCog, early cognition; EN, elastic net; PLS, partial least squares; TBSS, tract-based spatial statistics; CBSS, connectome-based spatial statistics.

**Figure 5.**
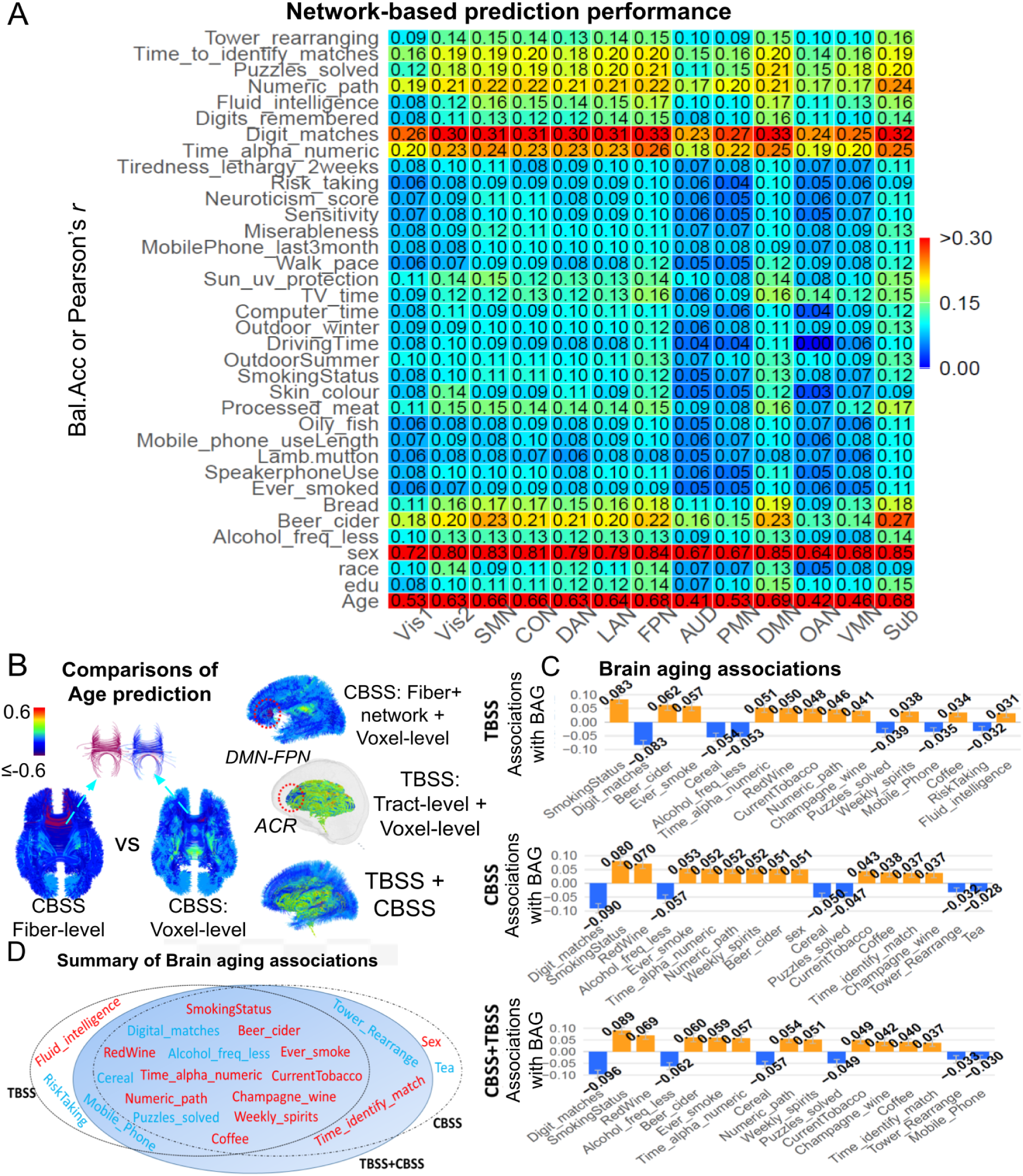
Network distributions of predictions and brain aging. **A.** Cognitive/behavioral predictions by using only pathways connecting each network based on the PLS model. Each cell reports balance accuracy or prediction correlation performance. **B.** Comparison of age-prediction contributions across CBSS fiber-level, voxel-level, CBSS (combined), TBSS, and combined TBSS+CBSS models. Contributions were quantified as the correlation between predicted age and FA features at the fiber or voxel level. **C.** Trait associations with the BAG for TBSS, CBSS and TBSS+CBSS approaches. **D.** Comparison of identified traits that are significantly associated with brain aging. Red and blue colors indicate negative and positive associations on brain aging. ACR, anterior corona radiata; BAG, brain age gap; PLS, partial least square regression model.

### Age prediction and brain aging analyses

We next compared the spatial signatures of age prediction across feature scales by evaluating prediction contributions at the fiber or voxel level, quantified as the correlation between predicted age and the corresponding FA feature at each location (**Figure 5B**, left). Voxel-level CBSS localized age-related contributions more precisely to tract subsegments than fiber-averaged models, revealing along-fiber heterogeneity that is obscured in fiber-level summaries, particularly in the genu of the corpus callosum linking left and right default mode regions, including L_9a, L_9m, L_9p and L_8BL, and R_9p, R_9a, R_9m and R_8BL. Although both CBSS and TBSS highlighted overlapping high-signal regions, CBSS linked significant effects to network-defined tractography connections, with particularly strong age sensitivity in DMN–FPN circuitry connecting L_10pp, L_p10p, L_10r and L_10d to R_10pp, R_p10p, R_a10p, R_OFC and R_10d, whereas TBSS localized signals at the same location to the WM tracts such as anterior corona radiata. We then quantified brain aging using the brain-age gap (BAG). Because raw BAG (predicted age − chronological age) remained correlated with chronological age, we applied the standard bias correction of *Smith et al.* ^29^, to derive an approximately age-independent deviation (**Supplementary Figure S14)**. We then used BAG as a surrogate marker of the extent of brain aging for downstream association analyses with behavioral, cognitive, and demographic phenotypes. Across methods, BAG showed systematic associations with behavioral and cognitive measures (**Figures 5C** and **5D**): smoking– and alcohol-related exposures were associated with older-appearing brains, whereas better cognitive performance was associated with younger-appearing brains, with Digit Match among the strongest negative associations. Although CBSS– and TBSS-derived age-prediction models showed broadly concordant overall association profiles (**Figure 5D**), CBSS enabled network-level aging analyses that localized smoking– and red-wine-related associations to distributed connectome systems, with particularly prominent effects in the aging of subcortical network (**Supplementary Figure S15**). TBSS captured similar global voxel-level trends, but could not directly identify network-level association patterns. Besides, TBSS additionally identified mobile phone use and risk-taking, but also showed an atypical positive association between BAG and fluid intelligence that is difficult to interpret biologically and may reflect residual confounding or model sensitivity. The combined TBSS + CBSS model yielded a more biologically coherent association profile, suggesting that the two separate representations may capture complementary aspects of brain aging.

### Lifespan growth charts

Across network pairs, FA-based SC lifespan trajectories from 3 to 90 years old show a robust inverted-U lifespan profile (**Figure 6A**), with increases through childhood and adolescence, a peak in early-to-mid adulthood, and a gradual decline thereafter. This pattern is most pronounced for sensory systems, including Vis1, Vis2, SMN, and their couplings, which tend to show earlier peaks and higher absolute FA, consistent with earlier maturation of primary sensorimotor pathways. By contrast, many higher-order association and control pairs, including DMN, FPN, CON, DAN and their cross-network edges, show more protracted growth and later peak ages, consistent with prolonged maturation of long-range integrative circuitry. Most peak ages fell between *18* and *45* years. Sex-stratified trajectories reveal age windows with significant sex differences, concentrated mainly around developmental and early-adult transitions and, for some edges, re-emerging in late adulthood aging. Where differences are present, annotated peak ages indicate earlier peaks in males than females for many network pairs. The observed patterns are broadly consistent with prior studies reporting inverted-U white-matter trajectories, tract-dependent peak timing, and later maturation in females for a majority of pathways ^30–32^.

**Figure 6.**
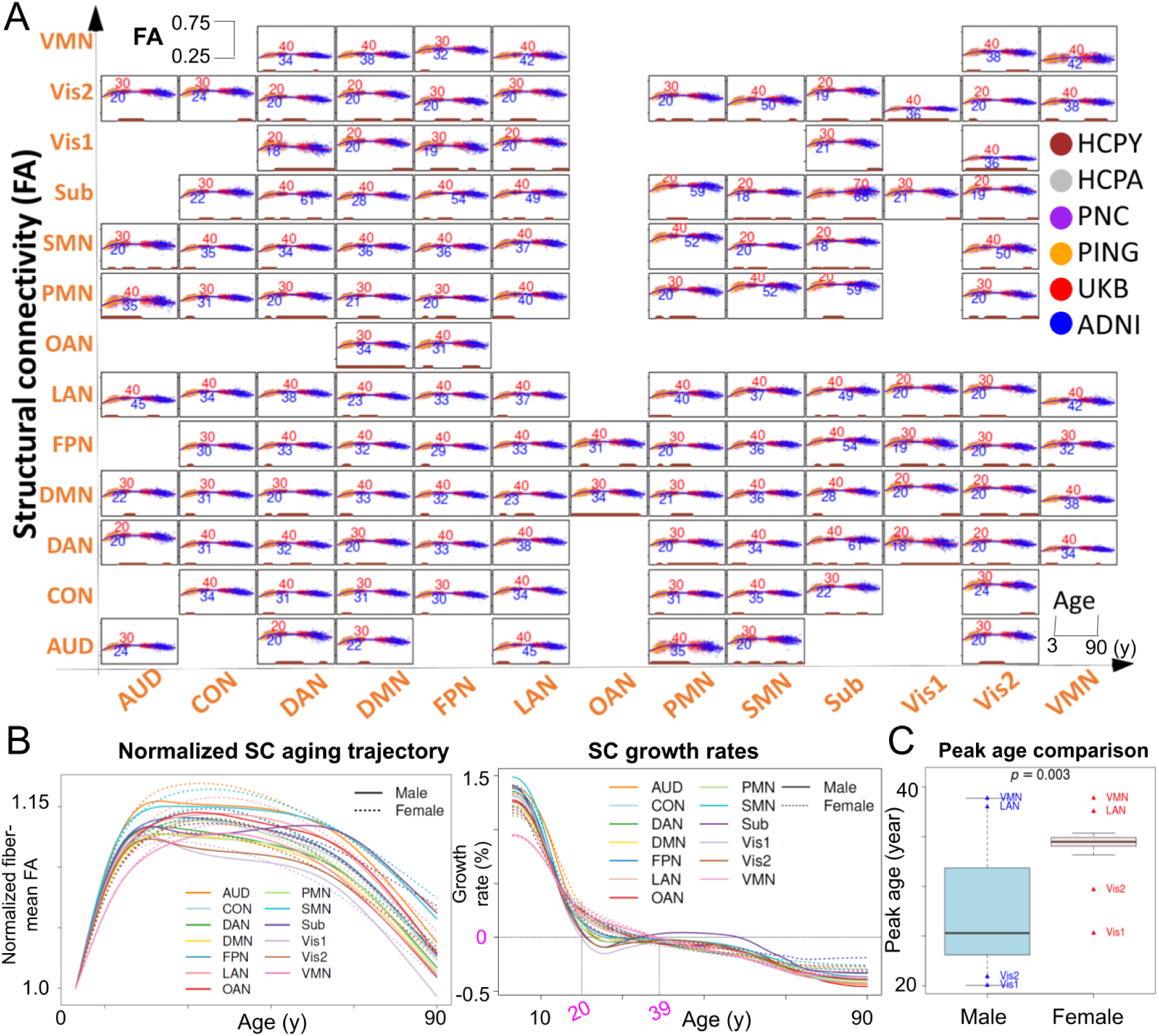
Brain structural connectivity growth chart. **A.** Lifespan growth curves for the mean brain structural connectivities are fitted between each network pair, with age spanning 3-90 years for males (blue) and females (red), separately, from six lifespan datasets (data points in six colors) across all network pairs. Peak ages were annotated (blue for male, and red for female); age ranges with sex difference were highlighted in brown segments at the x-axis. **B.** Normalized SC aging trajectories (left) and growth rates (right), averaged across all network pairs involving each network. Values are normalized to the fitted mean-FA at the age of three. Solid and dotted lines denote male and female trajectories, respectively. SC, structural connectivity; FA, fractional anisotropy. **C.** Boxplots for comparison of peak ages across 13 networks between females and males. The *p*-value was calculated based on the Wilcoxon signed-rank test. Peak ages for the four female outlier networks (Vis1, Vis2, VMN and LAN) are annotated, together with the corresponding peak ages for the same networks in males for comparison.

At the network-average level, the same developmental ordering was preserved (**Figure 6B**). Vis1, Vis2 and SMN peaked earliest and showed a relatively prolonged post-peak plateau, whereas higher-order association and control systems, including FPN, CON, LAN and VMN, peaked later and transitioned more directly into decline. Across networks, peak ages fell between *20* and *39* years, defining a general peak-to-stabilization window. Females showed later peak ages than males across all 13 networks at the network-average level (**Figure 6C**). The subcortical aggregate showed a more complex midlife profile, likely reflecting heterogeneity across constituent pathways rather than uniform continued maturation. Overall, these patterns are consistent with a sensorimotor-to-association gradient of white-matter development reported in prior lifespan diffusion MRI studies ^33^.

## Discussion

### Atlas-referenced multi-scale structural connectivity mapping

Large-scale diffusion-MRI efforts, including ENIGMA ^34^, UK Biobank and the lifespan Human Connectome Project ^13^, span over *100,000* participants, offering unprecedented opportunities to chart how WM networks mature, reorganize and decline across the human lifespan. Capitalizing on these data requires analytic tools that are computationally tractable, robust to inter-subject misalignment, reproducible across cohorts, and interpretable at the level of anatomical pathways and functional systems. Meeting all these criteria simultaneously has remained a major challenge in the field.

State-of-the-art whole tractography algorithms reconstruct pathways by propagating streamlines through local orientation fields derived from dMRI. The principle of inferring connectivity from local orientation fields involves the issues of pathways overlap, cross, branch, and have complex geometries ^35–37^. A previous study ^38^ investigating *96* distinct tractography pipelines reported that half of invalid bundles occurred systematically across *20* research groups. Computational power is another factor more and more demanding. Tractography-based connectome pipelines are substantially time-consuming: TractoFlow reported full diffusion and tractography processing times exceeding 5 hours per subject ^3^. Recent UK Biobank structural connectome studies have relied on computationally intensive pipelines, including multi-shell multi-tissue spherical deconvolution, anatomically constrained tractography with 1 million streamlines per subject, SIFT2 weighting and connectome construction at biobank scale ^39^. By contrast, registration and projection-based approaches are relatively efficient, relying mainly on registration and projection; for example, a typical 1-mm isotropic unimodal registration with MMORF takes about 5 minutes ^40^. Spatial alignment remains difficult in tractography, and current approaches address it mostly through dMRI image registration, ROI parcellation, and pathway clustering, while direct streamline registration is still emerging ^41^. Besides, WM fiber geometry varies markedly across individuals, scan quality differs across age ranges and protocols, and tractography introduces region– and method-specific biases ^38,42^. In comparison, registration and skeleton-based methods are relatively scalable and robust, but the WM microstructure is not inherently organized in connectome form. To bridge these regimes, we developed CBSS as an atlas-referenced framework that combines the efficiency of registration-and-projection workflows with the pathway– and network-level interpretability of structural connectomes. By projecting diffusion-derived measures from new cohorts onto the common space, CBSS avoids subject-level whole-brain tractography in downstream datasets and therefore reduces computational burden at biobank scale. At the same time, the atlas provides a common anatomical reference for cross-subject spatial correspondence, while local projection and along-path alignment help mitigate mismatch of white-matter geometry. Because the resulting traits are defined in pathway and network space, CBSS also supports direct analyses of connectome organization and structure–function relationships that are less naturally captured by voxelwise or tract-summary representations.

### Integrative multi-scale structural connectivity evaluation

A further strength of the present study is the matched evaluation framework used to interrogate structural connectome features across scales. Reproducibility and heritability provide one important test of whether these features capture stable and biologically meaningful variation rather than tractography-specific noise. Prior work ^43^ has shown that voxelwise diffusion metrics such as FA and MD are generally highly reproducible (ICC *> 0.8*), whereas tractography-derived networks are more variable and frequently require thresholding or harmonization to achieve stable estimates across scans or sites ^44^. Large-scale imaging genetics has likewise shown substantial genetic influence on white-matter microstructure, with moderate-to-high heritability (*0.3-0.7*) for tract-averaged diffusion traits ^21,45^, while tractography-derived connectivity measures remain heritable but more variable and highly polygenic (*0.03-0.3*) ^39^. Consistent with this literature, BFSC-derived traits in our study were both reproducible and heritable at scale. Structure–function coupling provides a second, complementary benchmark: anatomical connectivity constrains functional organization, and explicit structural edges offer a more direct substrate for SC–FC comparison than voxelwise or tract-summary measures alone ^27^. In our data, fiber-level CBSS recovered broad SC–FC correspondence, whereas voxel– and network-level embeddings further improved spatial correspondence and subject identifiability, indicating that voxel-level and hierarchical network representations preserve subject-specific information beyond streamline averaging. A fourth test is phenotypic relevance. Recent large-scale studies have shown that structural connectome measures can predict age, sex and cognitive traits, although performance is typically modest and depends on feature construction and sample size ^27,46^. In this context, the modest but reproducible gains we observed in cognitive and behavioral prediction, together with improved anatomical and network-level interpretability, suggest that CBSS adds useful structural information beyond conventional tract summaries. Finally, the lifespan analyses show that the same atlas-referenced framework can capture coherent developmental and aging trajectories across systems, with earlier maturation in sensory pathways and more protracted change in higher-order association circuitry, in line with recent lifespan connectome studies ^47^. Together, these analyses provide a common basis for understanding what atlas-referenced structural connectome measures capture at fiber, voxel and network scales, and where their practical limits lie in large multi-cohort studies.

### Limitations and future work

Several limitations remain. Atlas-referenced measures are still affected by registration quality, streamline reconstruction biases and reduced sensitivity for very short-range or anatomically complex pathways. This becomes more obvious for neighboring cortical pairs, for example, weak structural connectivity in the visual network can coexist with strong functional coupling because tractography near the grey–white boundary is difficult to recover robustly. The predictive gains observed here were also generally modest; their value lies in the combination of scalability, reproducibility, and pathway– and network-level interpretability across multiple analyses and cohorts. Looking forward, AI-based approach ^27^ may further improve atlas construction, feature representation and downstream prediction. CBSS also provides a scalable route to imaging genetics of pathway– and network-level white-matter traits, complementing existing voxelwise and tract-averaged analyses. More broadly, the evaluation framework introduced may help support more systematic comparison of diffusion MRI processing pipelines, atlases and embedding strategies across tasks, datasets and structural scales.

## Data availability

The data used in this work were obtained from six publicly available datasets: the UK Biobank (UKB) study, the Human Connectome Project Young Adult (HCP-Y), the Human Connectome Project Aging (HCP-A), the Pediatric Imaging, Neurocognition, and Genetics (PING) study, the Philadelphia Neurodevelopmental Cohort (PNC) study, and the Alzheimer’s Disease Neuroimaging Initiative (ADNI) study. HCP-Y data for this study are available for download from the Human Connectome Project (www.humanconnectome.org). Users must agree to data use terms for the HCP-Young Adult before being allowed access to the data and ConnectomeDB; details are provided at https://www.humanconnectome.org/study/hcp-young-adult/data-use-terms. Data from the HCP Aging studies can be downloaded as part of the HCP-Lifespan 2.0 release, distributed by the NIMH Data Archive (https://nda.nih.gov). See https://www.humanconnectome.org/study/hcp-lifespan-aging/data-releases for more information about data use terms. Data used in the preparation of this article were obtained from the Alzheimer’s Disease Neuroimaging Initiative (ADNI) database (adni.loni.usc.edu). The ADNI was launched in 2003 as a public-private partnership, led by Principal Investigator Michael W. Weiner, MD. For up-to-date information, see adni.loni.usc.edu. Access to the PING Data Resource is available through an online web interface at http://pingstudy.ucsd.edu. The PNC data is available at the dbgap with study accession phs000607.v3.p2.

## Code availability

We made use of publicly available software and tools. All codes used to generate results that are reported in this paper are available upon request.

## Acknowledgement

This work was partially supported by the National Institute on Aging (NIA) of the National Institutes of Health (NIH) grants [R01AG085581, RF1AG082938, RF1AG098697 to TF.L. and H.Z., and K01AG095286 to TF.L.], NIH grants [U01AG088667 to J.L.S., TF.L. and H.Z.; R01MH136055 to H.Z. and TF.L.; OT2OD038045 to H.Z.; and R21HD120911 to TF.L.]. The content is solely the responsibility of the authors and does not necessarily represent the official views of these institutes. We thank the individuals represented in the UK Biobank, ADNI, HCP-Y, HCP-A, PING and PNC datasets for their participation and the research teams for their work in collecting, processing and disseminating these datasets for analysis. This research has been conducted using the UK Biobank resource (application number 22783), subject to a data transfer agreement. We thank the individuals represented in the UKB study for their participation and the research teams for their work in collecting, processing and disseminating these datasets for analysis. We thank University of North Carolina at Chapel Hill and the Research Computing groups for providing computational resources and support that have contributed to the research results. Part of data collection and sharing for this project was funded by the Alzheimer’s Disease Neuroimaging initiative (ADNI) (National Institutes of Health Grant U01 AG024904) and DOD ADNI (Department of Defense award number W81XWH-12-2-0012). ADNI is funded by the National Institute on Aging, the National Institute of Biomedical Imaging and Bioengineering and through generous contributions from the following: Alzheimer’s Association; Alzheimer’s Drug Discovery Foundation; Araclon Biotech; BioClinica, Inc.; Biogen Idec Inc.; Bristol-Myers Squibb Company; Eisai Inc.; Elan Pharmaceuticals, Inc.; Eli Lilly and Company; EuroImmun; F. Hoffmann-La Roche Ltd and its affiliated company Genentech, Inc.; Fujirebio; GE Healthcare; IXICO Ltd; Janssen Alzheimer Immunotherapy Research & Development, LLC; Johnson & Johnson Pharmaceutical Research & Development LLC; Medpace, Inc.; Merck & Co., Inc.; Meso Scale Diagnostics, LLC; NeuroRx Research; Neurotrack Technologies; Novartis Pharmaceuticals Corporation; Pfizer Inc.; Piramal Imaging; Servier; Synarc Inc.; and Takeda Pharmaceutical Company. The Canadian Institutes of Health Research is providing funds to support ADNI clinical sites in Canada. Private sector contributions are facilitated by the Foundation for the National Institutes of Health (www.fnih.org). The grantee organization is the Northern California Institute for Research and Education, and the study is coordinated by the Alzheimer’s Disease Cooperative Study at the University of California, San Diego. ADNI data are disseminated by the Laboratory for Neuro Imaging at the University of Southern California. Part of the data collection and sharing for this project was funded by the Pediatric Imaging, Neurocognition and Genetics Study (PING) (U.S. National Institutes of Health Grant RC2DA029475 and R01HD061414). PING is funded by the National Institute on Drug Abuse and the Eunice Kennedy Shriver National Institute of Child Health & Human Development. PING data are disseminated by the PING Coordinating Center at the Center for Human Development, University of California, San Diego. Support for the collection of the PNC datasets was provided by grant RC2MH089983 awarded to Raquel Gur and RC2MH089924 awarded to Hakon Hakonarson. All PNC subjects were recruited through the Center for Applied Genomics at The Children’s Hospital in Philadelphia. HCP-Y data were provided by the Human Connectome Project, WU-Minn Consortium (Principal Investigators: David Van Essen and Kamil Ugurbil; 1U54MH091657) funded by the 16 NIH Institutes and Centers that support the NIH Blueprint for Neuroscience Research; and by the McDonnell Center for Systems Neuroscience at Washington University. HCP-Y data were supported by the National Institute On Aging of the National Institutes of Health under Award Number U01AG052564. The content is solely the responsibility of the authors and does not necessarily represent the official views of the National Institutes of Health.

## Authors’ contributions

TF.L., Z.Z. and HT.Z. proposed the concept and designed the methodology. HT.Z. supervised the project. TF.L., XF.W., and M.C. worked on the HCP-Y data to develop the pipeline. TF.L., Z.S., and Z.J. performed data analyses. TF.L., Z.S., Z.J., and X.Q. processed the MR imaging data for UK Biobank, HCP-A, PNC, PING, and ADNI datasets. TF.L. and S.G. generated the visualizations. TF.L. drafted the initial manuscript. Z.Z., M.D., J.L.S., XF.W., X.W.,TY.L., T.E.N., HP.Z., and HT.Z. suggested revision ideas and revised the manuscript. All authors critically reviewed the manuscript and approved the final version.

## Additional information

### Supplementary information

The online version contains supplementary materials.

**Correspondence and requests for materials** should be addressed to Hongtu Zhu.

## Online Methods

### CBSS workflow

We introduce CBSS, a comprehensive framework designed for fiber-based spatial statistical analysis in large-scale neuroimaging studies. CBSS leverages advanced tractography and structural connectome methodologies to effectively capture and represent microstructural fiber integrity. The framework consists of three major stages: (i) whole-brain tractography and alignment to standard space; (ii) construction of the BFSC atlas; and (iii) atlas-based mapping and quantification of fiber-, voxel-, and network-level spatial statistics (**Supplementary Figure 1**). Additional details are provided in the **Supplementary text**.

In the first stage, we construct a robust whole-brain structural connectome. This involves generating high-quality whole-brain tractographic data derived from diffusion MRI (dMRI) and precisely mapping structural connections. Specifically, we processed the dMRIs and generated whole-brain tractography for each subject using TractoFlow ^3^ and anchored streamline endpoints to a fine-grained parcellation derived from the HCP-MMP cortical atlas ^14^ and FreeSurfer subcortical labels to define the nodes of the SC network. To improve parcel-level stability, we used the 43 pairs of HCP test–retest scans to identify cortical parcels for merging based on the following selection criteria: they were spatially adjacent, belonged to the same functional network, had mean correlation greater than 0.7 between their fiber-count profiles with other ROIs, and would not reduce the fiber count test-retest reliability after merging. Applying these criteria, 26 cortical parcels were merged into 12 larger regions (**Supplementary Figure S2**). Together with 13 subcortical regions, this yielded a final parcellation of 359 regions. Following Cole-Anticevic Brain-wide Network Partition (CAB-NP) ^15^, these 359 parcels are labeled as 12 canonical functional networks plus a subcortical network. To reduce incomplete and false-positive connections, we further applied cortex surface dilation, streamline cutting and outlier streamline removal, following the PSC strategy ^48^. We then map the cortical and subcortical ROI labels to the points on individual cortical surfaces using surface registration. Since all the generated fiber curves connect cortical regions intersect with cortical surfaces, we extract the endpoints of the fibers and snap the endpoints of each fiber to the nearest points on the individual cortical surface by minimizing the L₂ distance such that we can assign the ROI labels of the points on cortical surface to the fiber endpoints ^49^. (iii) We group fibers based on the connections of their endpoints with different ROIs after parcellation and we calculate the fiber counts between different ROI pairs for each subject. The output of Stage 1 is a parcellation of fibers *F_it_* into 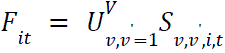 where 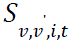 is the tractography that connects the ROI pair *R_v_* and *R_v’_* for subject *i* at visit age *t*.

The second stage focuses on generating a reliable parcellation-based fiber skeleton, ensuring data integrity and statistical robustness. Initially, pre-clustering quality control procedures remove outliers by filtering fibers with fiber count reliability (ICC) less than *0.4*, and merging *14* ROIs from the Glasser 360 atlas that are physically neighboring, from the same network, exhibit highly similar connectivity patterns (correlations of fiber counts *> 0.7*) and guarantee the merging did not decrease the overall reliability of fiber counts. Clustering methods then identify core structural fiber pathways by grouping fibers connecting selected ROI pairs (**Supplementary Figure S16A**). We utilize the Tract Dictionary Learning (TractDL) method ^16^, to cluster the fibers into *6,090* clusters (**Supplementary Figure S16C**). After clustering fibers between all ROI pairs, we extract the centroid fiber from each cluster. The centroid of a cluster 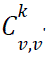 is defined as the fiber with the smallest median MDF distance from all other fibers within the cluster. Specifically, the centroid fiber of each cluster is defined as:

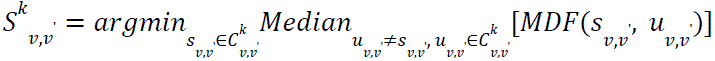

Post-clustering quality control further refines the clusters by eliminating small clusters and centroid fibers with insufficient FA test-retest reproducibility (FA *< 0.4*). The final output of this step is the raw fiber skeleton atlas composing of all cluster centroid fibers:

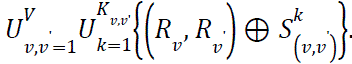

In the third stage, fiber-based spatial statistics from multiple life-span neuroimaging datasets are accurately mapped onto the developed fiber skeleton. This step first involves projecting individual-level fiber statistics, such as fractional anisotropy (FA), onto the skeleton. Then, elastic registration techniques are employed to align individual fiber skeletons across subjects while removing statistical outliers, thereby facilitating group-level comparisons. Sparse representation methods are utilized to efficiently represent tractographic data, enhancing interpretability and computational efficiency.

The third stage maps individual spatial statistics onto the atlas through two steps: projection of fiber-based statistics and elastic registration. First, voxel-based spatial metrics—such as FA and MD—are projected onto the refined fiber skeleton using a modified TBSS approach. FA images are first nonlinearly registered to the 1×1×1 mm³ MNI152 space. For each voxel on the atlas skeleton, the projection is made by searching perpendicularly to the local tract direction for the maximum FA value in the subject’s image. This voxel is recorded as the tract’s center, and its coordinates are used to extract corresponding values for all other spatial statistics. Next, we use a finer 1×1×1 voxel neighborhood to preserve tract detail, combined with 0.1 mm³ trilinear interpolation for sub-voxel precision. The result is a set of subject-specific tract profiles containing projected statistics, enabling alignment-invariant and population-level tract analysis. The second stage is the elastic registration. Despite nonlinear registration of diffusion measures to MNI space in prior steps, residual fiber-level misalignment can occur because corresponding along-tract profiles may still differ by local stretching or compression. To reduce this along-fiber variability, we performed cross-subject elastic registration of streamline-wise diffusion profiles using the Fisher–Rao/square-root slope function framework, which separates phase from amplitude variation and aligns each subject-specific profile to a cross-subject Karcher mean template ^50^ (**Supplementary text**; **Supplementary Figure S16F**). We estimated a smooth monotone warping function for each streamline profile and regularized the warp toward the identity so that alignment corrected local timing or position differences without introducing excessive distortion; the penalty was additionally scaled by streamline length. This procedure yields reparameterization-invariant correspondence of diffusion profiles across subjects and produces aligned tract-based diffusion features for downstream population-level graph analyses^51^.

### UK Biobank SNP data and heritability analyses

We conducted a GWAS of CBSS fiber-level FA features using dMRI data and the imputed genetic variants data from UK biobank phase 1-5 all white individuals. We further performed the following genetic variants data quality controls on subjects with both imaging and genetics data in each study: 1) excluded subjects with more than *10%* missing genotypes; 2) excluded variants with minor allele frequency less than *0.01*; 3) excluded variants with missing genotype rate larger than *10%*; 4) excluded variants that failed the Hardy-Weinberg test at *1e-7* level; and 5) removed variants with imputation INFO score less than *0.8*. Then we estimate the SNP heritability for each fiber-level phenotype by all autosomal variants using GCTA-GREML analysis ^52^. The adjusted covariates included age at imaging, age-squared, sex, age-sex interaction, age-squared-sex interaction, imaging site, and the top *40* genetic principal components (PCs) provided by UKB (Data-Field 22009). The heritability estimates were tested in one-sided likelihood ratio tests.

### Charting lifespan aging trajectories on tractography

After tractography was finalized, the mean FA across fibers connecting each network pair was modeled using generalized additive models (GAMs) ^53^ to estimate the lifespan trajectories of SC between networks. This analysis integrated UKB, HCP-YA, HCP-A, PNC, PING, and cognitively normal baseline participants from ADNI-GO/2. For each feature, we fit 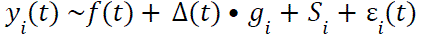 where *t* is age, and *g_i_* is sex (*1* for male and *0* for female), and *S_i_* is the study site for subject *i* at time *t*. The smooth terms *f*(*t*) and Δ(*t*) were modeled with cubic regression splines, representing the overall lifespan trajectory and the age-dependent sex effect, respectively, and 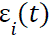 denotes Gaussian noise. Sex-specific trajectories were estimated from the fitted GAM using the mgcv package in R (v4.4.0), with smoothing parameters selected by restricted maximum likelihood (REML). Peak ages were defined as the maxima of the fitted curves for male and female, respectively. Age ranges with significant sex-difference intervals were determined from the age-varying male–female difference estimated by the GAM, and were defined as age ranges where the *95%* simultaneous confidence interval excluded zero after correction for multiple testing of *66* network pairs.

### Cognitive and behavioral prediction in lifespan datasets

We evaluated phenotype prediction using CBSS and TBSS features in UKB, HCP-YA, HCP-A, PNC, PING and ADNI-GO/2. FThe UKB data field, data type and explanation of *96* cognitive, behavioral and demographic outcomes we predicted are listed in **Supplementary Table 1**. Similar to Jamison, *et al.* ^27^, for each dataset except UKB, participants with complete phenotype and imaging data were partitioned into age-stratified training, validation and test sets with approximate proportions of 7:1.5:1.5. UKB training, validation and test data were split following **Figure 3A**. To reduce information leakage from related individuals, data splits were constructed at the family level in HCP using parental information, and at the participant level in ADNI to account for repeated longitudinal entries, such that members of the same family or multiple scans from the same participant were always assigned to the same split. All models were trained on the training data, with hyperparameters tuned on the validation data, and the final model performance was assessed in the untouched test subset.

PC embeddings were extracted and feature matrices were constructed separately for CBSS fiber-level, voxel-level, network-level, and TBSS voxel-level and combined CBSS + TBSS representations; all predictors were z-scored before model fitting. For TBSS, we retained up to 200 PCs together with the mean for each white-matter tract block, capturing more than 99% of the variance. For CBSS voxel-level features, we retained the first three principal components (PCs) and the mean for each fiber; the first three PCs explained more than 50% of the variance for over 90% of fibers. For CBSS network-level features, we further constructed hierarchical PCs from the pooled top three PCs of all constituent fibers within each network pair, and the number of retained hierarchical PCs was determined by a validation-tuned cumulative contribution threshold. Continuous and ordinal outcomes were standardized before modelling. Multicategory outcomes were binarized against the majority class to maintain a common prediction framework; for example, race was coded as white versus non-white. PLS and EN predictive models were fitted, with hyperparameters including the number of PLS components (*1* to *30*), the elastic-net penalty parameters λ and α, the cumulative PC contribution threshold (*0.70*–*0.95*) applied separately to each network pair, and, for sex prediction, the probability threshold used to convert predicted scores into class labels. Tuning parameters were selected by maximizing area under the receiver operating characteristic curve (AUC) or prediction correlation (Pearson’s *r*) on the validation set. Final performance was reported as balanced accuracy for sex and as prediction correlation for all other outcomes, to provide a unified basis for comparison across phenotypes. For both the PLS and EN models, bootstrap confidence intervals for prediction performance were estimated using 100 bootstrap resamples of the training and test participants. Because model estimates were less stable in the smaller datasets, we additionally applied bagging within each bootstrap replicate for all datasets other than UKB, by fitting 20 models to resampled training data and averaging their predictions before evaluation on each of the 100 resampled test sets.

### SC-FC prediction

Whole-brain 64,620 ROI pairwise FC was predicted from SC features by first encoding FC into network-pairwise PC embeddings. Specifically, for each ROI pairwise FC, extreme values were identified using a 5× median absolute deviation (MAD) criterion and excluded, and FC vectors were z-scored across subjects to ensure scale-invariant comparison. Then for each of the 78 functional network pairs, we extracted PCs from the ROI pairwise between ROI pairs within the network pair, and SC-based prediction models were trained to predict the corresponding subject-level PC scores of FC based on CBSS fiber-level, voxel-level, network-level or TBSS features. Hyperparameters were selected on the UKB validation set (**Figure 3A**), after which the optimized model was applied to held-out participants. The predicted FC PC scores were then decoded to reconstruct subject-specific FC values for all *64,620* ROI pairs. We performed the prediction based on the PLS model, where the number of latent components (1 to 30) was selected on the validation set. For SC–FC prediction, TBSS voxel-level, CBSS voxel-level and CBSS network-level PC embeddings were constructed analogously to those used in behavioural and cognitive prediction, and the number of retained hierarchical PCs was determined by optimizing the prediction accuracy on the validation data. Prediction accuracy was evaluated by the Pearson correlation between predicted and observed FC profiles across ROI pairs within each subject, providing a global measure of connectome reconstruction. We then calculate the *avgcorr* and *avgrank* to evaluate the consistency between the predicted and observed FC ^27^:

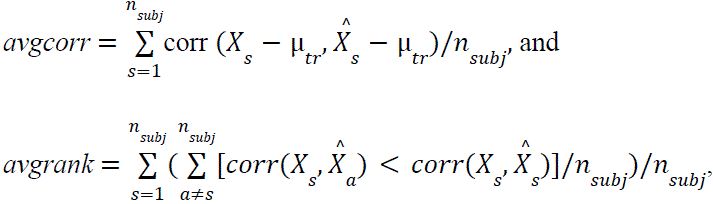

where 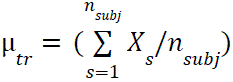 for training subjects, *n*_*subj*_ is the sample size and 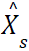 and *X_s_* incidate the predicted and observed functional connectivity for a subject *s*.

### Brain aging association analyses

For each structural representation, we computed BAG separately: using PLS-based predictions for CBSS fiber-level, voxel-level, network-level and TBSS voxel-level models, and using ensemble predictions from PLS and EN for the combined CBSS (fiber + voxel + network), combined TBSS (tract + voxel) and combined CBSS+TBSS models. We first generated age predictions in the UK Biobank phase-3 hold-out set using models trained on UK Biobank phase-1/2 data. Predicted age was back-transformed to years using the mean and standard deviation of chronological age in the training set, and BAG was defined as predicted age minus chronological age. Because raw BAG retained residual age dependence, we applied an out-of-sample linear age-bias correction ^29^: the hold-out set was randomly split into two halves, and in each half raw BAG was regressed on chronological age using the other half; the fitted age effect was then subtracted to yield an approximately age-independent corrected BAG. This cross-fitted corrected BAG was used in all downstream analyses. Associations between corrected BAG and *96* behavioural, cognitive and demographic traits were then tested separately for each imaging representation using linear regression, with standardized trait values entered one at a time as the predictor of interest. All models adjusted for imaging site, sex, quadratic age terms and sex-by-age interactions, brain-position covariates, volumetric scaling, race and education; when sex, race or education was the trait of interest, the corresponding covariate was omitted from the adjustment set. Statistical significance across traits was controlled by Bonferroni correction for 96 phenotypes.

