## Supplementary materials for "Connectome-based spatial statistics enabling large-scale population analyses of human connectome across cohorts"

**The supplementary materials include:**

1. Supplementary Text
2. Supplementary Figures S1-S17
3. Supplementary Table S1

### Supplementary text

Image acquisition

This work used diffusion magnetic resonance imaging (dMRI) data from six cohorts with distinct imaging protocols: Human Connectome Project Young Adult (HCP-Y), Human Connectome Project Aging (HCP-A), UK Biobank (UKB), the Philadelphia Neurodevelopmental Cohort (PNC), the Pediatric Imaging, Neurocognition, and Genetics study (PING), and the Alzheimer's Disease Neuroimaging Initiative (ADNI). Specifically, the image acquisition and preprocessing procedures were detailed in the UKB brain imaging documentation for the UKB study (https://biobank.ctsu.ox.ac.uk/crystal/crystal/docs/brain_mri.pdf), (1) for the HCP-Y study, (2) for the HCP-A study, (3) for the PNC study, (4) for the PING study, and (5) for the ADNI study. All participants provided informed consent, and each study received approval from the appropriate research ethics committees or institutional review boards. UKB was approved by the North West Multicentre Research Ethics Committee (11/NW/0382). PNC was approved by the institutional review boards of the University of Pennsylvania and the Children's Hospital of Philadelphia. PING was approved by the human research protection programmes and institutional review boards at the nine participating institutions. HCP was approved by the institutional review board at Washington University (201204036). HCP-A was approved by the Institutional Review Board at Washington University in St. Louis. The ADNI (ADNI GO and ADNI 2) studies were approved by the Institutional Review Board or Research Ethics Board at each participating site. . Informed written consent was obtained from all participants at each site.

**UKB data acquisition**

The scanner is a standard Siemens Skyra 3T running VD13A SP4, with a standard Siemens 32-channel RF receive head coil. Diffusion MRI in UKB was acquired at 2 × 2 × 2 mm spatial resolution using anterior-to-posterior phase encoding and a multiband acceleration factor of 3. The protocol included two non-zero b-value shells ($b = 1000$ and $2000 s/mm^2$), each with 50 diffusion-encoding directions. Diffusion preparation used a standard monopolar Stejskal–Tanner pulse sequence, with echo time (TE) of 92 ms and repetition time (TR) of 3,600 ms. Relative to a twice-refocused bipolar sequence, this design provides higher signal-to-noise ratio through the shorter TE, at the cost of stronger eddy-current distortion (6). Additional details are provided in Section 2.8 of the UKB brain imaging documentation.

**HCP-Y and HCP-A data acquisition**

Diffusion MRI in the HCP-Y cohort was acquired at 1.25 mm isotropic resolution in six runs (each approximately 9 min 50 s), spanning three diffusion gradient tables, each collected with both right-to-left and left-to-right phase-encoding directions. Each run included approximately 90 diffusion-weighted directions and six interspersed $b = 0$ images across three shells ($b = 1,000, 2,000$ and $3,000$ s/$mm^2$), with approximately balanced sampling across shells. For HCP-A, we used the two runs from 98 direction data with both AP and PA phase encoding directions, including 92 $b = 1500 s/mm^2$, 92 $b = 3000 s/mm^2$ and 14 $b = 0 s/mm^2$. The in-plane resolution was 1.5 mm. Additional acquisition details for HCP-Y and HCP-A have been described previously (7) and (8), respectively. In the BFSC atlas construction procedure using HCP-Y data, all shells were used for fODF reconstruction, whereas only the $b = 1,000$ s/$mm^2$ shell for HCP-Y and $b = 1,500$ s/$mm^2$ shell for HCP-A was used to derive DTI metrics. Of the 1,206 participants in the HCP-Y data, 1,042 with both raw diffusion MRI and T1-weighted MRI were included. Among these, 44 underwent repeat imaging approximately two weeks later, yielding 88 scans for test–retest analyses.

**PING, PNC and ADNI data acquisition**

For the PING data, diffusion MRI in PING was acquired across multiple 3 T Siemens, GE and Philips scanners at $2.5mm$ isotropic resolution using a single-shell protocol with $b = 1000s/mm^2$. The acquisition included 30 diffusion-encoding directions on GE and Siemens platforms and 32 directions on Philips platforms, together with two to four of $b = 0$ images. Additional details regarding PING image acquisition and preprocessing can be found in (4, 9). For the Philadelphia Neurodevelopmental Cohort (PNC) dataset, MRI data were acquired on 3T Siemens scanners using a standardized neurodevelopmental imaging protocol. The diffusion weighted imaging (DWI) sequence consisted of 64 diffusion-weighted directions with single shell $b = 1000s/mm^2$, 7 scans with $b = 0s/mm^2$, and a voxel resolution of approximately $2mm$ isotropic. For the ADNI, we used the ADNI-2/GO datasets. Diffusion MRI data were acquired on a 3.0 T GE scanner using the ADNI standard protocol, with a spatial resolution of $1.37 \times 1.37 \times 2.7mm^3$ spatial resolution. The majority of its scans included five b0 images and 41 diffusion-weighted acquired at $b = 1000s/mm^2$. More details regarding PNC and ADNI image acquisition and preprocessing can be found in (3, 10), respectively.

Standard dMRI data processing

Diffusion MRI data from UK Biobank and the smaller cohorts were preprocessed using cohort-specific diffusion MRI pipelines that were broadly consistent with standard FSL/TBSS workflows (11). Preprocessing included susceptibility-induced distortion correction when dedicated correction scans (e.g., reverse phase-encoding) were available, followed by eddy-current correction, motion correction, brain masking, and diffusion tensor fitting. In HCP-Y and HCP-A, distortion correction used paired diffusion acquisitions with opposing phase-encoding directions, whereas in UK Biobank it used a smaller number of reverse phase-encoding b0 images. The $b \leq 1500$ shells for HCP-A and $b \leq 1000$ shells for other dataset were fed into the diffusion-tensor-imaging (DTI) fitting tool DTIFIT, creating DTI outputs such as fractional anisotropy (FA), mean diffusivity (MD) and tensor mode (MO). The resulting diffusion maps were then nonlinearly registered to the FMRIB58 $1mm$ FA template in MNI152 standard space.

Detailed data processing of the CBSS

We developed the connectome-based spatial statistics (CBSS) framework in three main stages (**Supplementary Figure 1**): (i) individual whole-brain tractography of the HCP-Y data with alignment to standard space; (ii) construction of a brain function–aware structural connectome (BFSC) atlas based on aggregated whole-brain tractography in standard space; and (iii) mapping of other datasets onto the atlas and quantifying fiber-level, voxel-level, and network-level spatial statistics. In the following sections, we describe each module in detail. Stage one comprises three substeps: (S1.1) construction of high-quality individual whole-brain tractographic data; (S1.2) brain parcellation; and (S1.3) nonlinear registration to standard space.

**Individual Whole-brain tractography**

For each HCP-YA scan, whole-brain tractography was reconstructed using the state-of-the-art diffusion MRI pipeline *Tractoflow* (12), and tractography near the cortical boundary was further refined by incorporating cortical surface information through surface-enhanced tractography (SET) (13). The Tractoflow pipeline first preprocesses the diffusion-weighted images (DWIs), including denoising, susceptibility, eddy-current and motion correction, brain extraction, N4 bias-field correction, cropping, intensity normalization and resampling. It then computes diffusion tensor imaging (DTI) metrics and fiber orientation distribution function (fODF) metrics (14, 15).

Next, the T1-weighted image is processed through denoising, N4 bias-field correction, resampling, brain extraction and cropping, followed by registration to diffusion space, tissue segmentation, and construction of tracking masks and seeding maps. Finally, whole-brain tractography is generated using the probabilistic tracking algorithm of (16), based on the fODF image and seeding mask. To improve the anatomical precision of streamline terminations near cortex, we applied SET (13), which constrains streamline endpoints to intersect the cortical surface. This step reduces premature termination of streamlines within white matter and enables direct integration with cortical parcellations for connectome construction. We excluded streamlines with fewer than three points or with only one endpoint intersecting the cortical surface, and further retained only streamlines with lengths between 10 and 250 mm. To reduce incomplete and false-positive connections, we additionally applied cortical surface dilation, streamline cutting and outlier streamline removal, following the PSC strategy (17).

The output of step $(\mathbf{S1.1})$ for subject $i$ at time $t$ consists of a whole-brain tractography dataset, denoted by $\mathcal{F}_{it}$, together with associated diffusion profiles, denoted by $\mathcal{P}_{it}$ (for example, fractional anisotropy, FA). Specifically, $\mathcal{F}_{it} = \{F_{it,1}(\cdot), \ldots, F_{it,M_{it}}(\cdot)\}$, where $\mathcal{F}_{it}$ contains $M_{it}$ streamlines, and the $k$-th streamline $F_{it,k}$ is represented as an ordered sequence of $m_{(it,k)}$ 3D points $p_{(it,k),j} = \big(x_{(it,k),j},\, y_{(it,k),j},\, z_{(it,k),j}\big) \in \mathbb{R}^3, j = 1, \ldots, m_{(it,k)},\ k = 1, \ldots, M_{it}$. The corresponding diffusion-profile set is $\mathcal{P}_{it} = \{P_{it,1}(\cdot), \ldots, P_{it,M_{it}}(\cdot)\}$, where each $P_{it,k}(s)$ is a one-dimensional function sampled along the streamline $F_{it,k}(s)$. In the HCP-Y data, this procedure yielded approximately $M_{it} \approx 1$ million streamlines per individual. Tractography was generated with a step size of 0.2 mm, while in the downstream analysis, each streamline $F_{it,k}(s)$ was represented by 100 sampled points along the curve $m_{(it,k)} = 100$.

**Brain parcellation**

In $(\mathbf{S1.2})$, we generated an individual-subject 373-region parcellation for connectome construction. Cortical nodes were defined using the Glasser 360 atlas (HCP-MMP1.0) (18), which partitions the cortex into 360 parcels. This parcellation was used to define structural connectome nodes and to assign streamlines to pairs of regions of interest (ROIs) for subsequent fiber-count extraction. Each subject's cortical surface was reconstructed with FreeSurfer, and the HCP-MMP1.0 atlas was mapped onto the individual surface in surface space, yielding 360 subject-specific cortical ROIs. The cortical parcels were further assigned to 12 functional networks according to the Cole–Anticevic Brain-wide Network Partition (CAB-NP) (19), including including the visual 1 and 2 (Vis1 and Vis2), language network (LAN), default mode network (DMN), auditory network (AUD), somatomotor network (SMN), dorsal attention network (DAN), frontoparietal network (FPN), cingulo opercular network (CON), posterior multimodal network (PMN), ventral multimodal network (VMN) and orbito affective network (OAN). The Glasser360-CAB-NP parcellation improves neuroanatomical precision for studying the structural and functional organization of human brain. We additionally incorporated 13 FreeSurfer-derived subcortical ROIs from the subject's anatomical segmentation which we referred to as a subcortical network (Sub), including the brainstem and bilateral thalamus proper, caudate, putamen, pallidum, hippocampus and amygdala. Together, these procedures yielded 373 subject-specific ROIs (360 cortical and 13 subcortical) for SC extraction across subjects.

**Registration and extraction of ROI-level connectivity**

In $(\mathbf{S1.3})$, we aligned subject-specific tractography to a common MNI152 reference space and extracted ROI-wise structural connectivity. An MNI152 cortical ribbon was first generated by processing the FSL MNI152 1 mm template with FreeSurfer `recon-all` (20). Individual cortical surfaces were then registered to this common surface using FreeSurfer surface registration

(21), and the resulting surface-to-volume transformations were propagated to the tractographic data. Cortical ROI labels were transferred to the individual cortical surface through this registration, whereas subcortical ROIs were retained in volume space. Because streamline endpoints were constrained to intersect the cortical surface, each cortical endpoint was assigned the label of its nearest surface vertex, and subcortical endpoints were labeled according to the volumetric ROI they intersected. Streamlines were then grouped by the ROI pairs based on labels of their two endpoints, and fiber counts were computed for each ROI pair in each subject. This yielded a partition of $\mathcal{F}_{it}$ as $\mathcal{F}_{it} = \bigcup_{v,v'=1}^{V} \mathcal{S}_{v,v',i,t}$, where $\mathcal{S}_{v,v',i,t}$ denotes the set of fibers connecting ROIs $R_v$ and $R_{v'}$, and $V = 373$ denotes the initial number of nodes before later ROI merging, including 360 cortical and 13 subcortical regions. Next we will list detailed substeps in stage two: (S2.1) preclustering QC for outlier filtering; (S2.2) preclustering QC for ROI merging and filtering; (S2.3) fiber clustering and atlas construction; and (S2.4) post-clustering QC.

**Pre-clustering QC: outlier filtering.**

In $(\mathbf{S2.1})$, for each subject, we performed the pre-clustering quality control (QC) by removing outlier streamlines that did not follow major white-matter (WM) pathways, in order to improve bundle definition and stabilize ROI-pairwise structural connectivity estimates. Such outliers are common in tractography because of the limited spatial resolution of diffusion MRI and the sensitivity of streamline reconstruction to local modeling and tracking parameters (22). Following (23), we used QuickBundles with the minimum average direct-flip (MDF) distance (24, 25) to cluster streamlines and detect small, isolated clusters as putative outliers. Based on the MDF distance, QuickBundles iteratively grouped streamlines by assigning each streamline to an existing cluster when its distance to the cluster centroid was smaller than a prespecified threshold $\theta_t$; otherwise, a new cluster was created; and cluster centroids were updated in both cases. Finally, Clusters with very small membership were treated as outlier clusters and removed. Representative outlying streamlines are shown in red in **Supplementary Figure 17** for one selected ROI pair. During this procedure, for each streamline $s = [s_1, s_2, \ldots, s_K]$ and its flipped representation $s^F = [s_K, s_{K-1}, \ldots, s_1]$, the MDF distance between $s$ and another streamline $u$ is defined as

$$d_{\text{direct}}(s,u) = \frac{1}{K}\sum_{i=1}^{K} \|s_i - u_i\|, d_{\text{flipped}}(s,u) = d(s,u^F) = d(s^F,u),$$

$$\text{MDF}(s,u) = \min\left\{d_{\text{direct}}(s,u),\ d_{\text{flipped}}(s,u)\right\},$$

where $\|\cdot\|$ denotes the $\mathbb{L}_2$ distance. MDF distance is very fast and easy to compute and also considers the fiber orientation issues. After calculating the MDF distance of each pair of the fibers, the clustering process, similar to $k$-means (26), is performed. In the HCP-YA data, the clustering threshold $\theta_t$ strongly affected the number of detected outliers. Consistent with our previous findings (23), thresholds $\theta_t > 10$ mm removed few apparent outliers, whereas thresholds $\theta_t < 5$ mm were overly stringent. In earlier work, we used $\theta_t = 8$ mm to remove obvious outlying streamlines. Here, to obtain a cleaner tractography skeleton for subsequent atlas construction, we used a more stringent threshold of $\theta_t = 5$ mm. The output of $(\mathbf{S2.1})$ was an outlier-filtered tractography set for each scan, $\mathcal{F}_{it}^{(1)} = \bigcup_{v,v'=1}^{V} \mathcal{S}_{v,v',i,t}^{(1)}$, where $\mathcal{S}_{v,v',i,t}^{(1)}$ denotes the set of streamlines connecting $R_v$ and $R_{v'}$.

**Pre-clustering QC: merging ROIs and filtering ROI pairs via fiber count**

In $(\mathbf{S2.2})$, we merged ROIs with similar connectivity patterns to reduce the degree of freedom and increase reproducibility. Let $R_v^{(0)}$ and $\mathcal{S}_{v,v,i,t}^{(0)}$ denote the ROI parcellation and ROI pairwise fiber set before ROI merging. Specifically, for ROI $R_v^{(0)}$ of subject $i$ at time $t$, we extract its fiber count vector with the other ROIs and obtain the fiber count vector $N_{v,i,t} = (\|\mathcal{S}^{(0)}_{v,v',i,t}\|_0)^V_{v'=1}$, where $\|\mathcal{S}\|_0$ denotes the number of a set $\mathcal{S}$. Afterwards we calculate the averaged correlation of the fiber count vector between ROIs $R^{(0)}_v$ and $R^{(0)}_{v'}$ across all the subjects $C_{v,v'} = \sum_{i=1}^n \text{Cor}(N_{v,i,t_i}, N_{v',i,t_i})/n$. We merge fiber tracks in $R^{(0)}_v$ and $R^{(0)}_{v'}$ together, if $R^{(0)}_v$ and $R^{(0)}_{v'}$ are the ROIs which have large mean correlation $C_{v,v'} > 0.7$, belong to the same functional network and are spatially neighboring, and the merging would not significantly reduce the test-retest reliablity. **Supplementary Figure S16A** shows the distributions of ICC values before and after ROI merging. ICCs were slightly higher after merging, with a smoother distribution shifted toward larger values, indicating improved stability. In this way, we merge ROIs while preserving the functional network structure. These fiber curves and their endpoints are merged accordingly to refine the initial parcellation. The output is the refined $\widetilde{V}$ ROIs $R^{(1)}_v, v = 1, \cdots, \widetilde{V}$ after merging, and a set of fiber bundles across all the merged ROI pairs for each scan $\mathcal{F}^{(1)}_{it} = \bigcup_{v,v'=1}^{\widetilde{V}} \mathcal{S}^{(1)}_{v,v',i,t}$, where $\mathcal{S}^{(1)}_{v,v',i,t}$ is the tractograph of fibers between the updated ROIs $R^{(1)}_v$ and $R^{(1)}_{v'}$. Next, we filter out ROI pairs and related fibers with low reproducible fiber counts. In particular, we quantify the reproducibility of ROI fiber counts on the HCP test-retest dataset by using the intraclass correlation coefficient (ICC) (27, 28) at the test-retest dataset, which is defined as

$$\text{ICC}_{v,v'} = \frac{1}{ns^2_{v,v'}} \sum_{i=1}^n \left(N_{(v,v'),i,t_{i,1}} - \bar{N}_{v,v'}\right) \left(N_{(v,v'),i,t_{i,2}} - \bar{N}_{v,v'}\right)$$

where $N^{(1)}_{(v,v'),i,t_{i,j}} = \|\mathcal{S}^{(1)}_{v,v',i,t_{i,j}}\|_0$ is the fiber number connecting $R^{(1)}_v$ and $R^{(1)}_{v'}$ in $\mathcal{S}^{(1)}_{v,v',i,t}$ for the $i$-th subject at baseline $t = t_{i,1}$ and at retest time $t = t_{i,2}$ respectively; $\bar{N}_{v,v'}$ and $s^2_{v,v'}$ are the mean and standard deviation of fiber counts in the whole test-retest dataset $\bar{N}_{v,v'} = \sum_{i=1}^n [N^{(1)}_{(v,v'),i,t_{i,1}} + N^{(1)}_{(v,v'),i,t_{i,2}}]/(2n)$ and $s^2_{v,v'} = \sum_{i=1}^n \left[\left(N^{(1)}_{(v,v'),i,t_{i,1}} - \bar{N}_{v,v'}\right)^2 + \left(N^{(1)}_{(v,v'),i,t_{i,2}} - \bar{N}_{v,v'}\right)^2\right]/(2n-1)$. We discard the fiber curves in the ROI pair with low $\text{ICC}_{v,v'}$ (**Supplementary Figure S16B**). Specifically, we recalculate ICCs on the merged ROIs. An ICC of less than $0.4$ usually represents poor reproducibility(29), and hence we discard such ROI pairs. We also remove the ROIs with the percentage of the missingness at the test-retest dataset larger than $50\%$ to ensure the connectivity between any two ROIs exist for the majority of the subjects. The output of (S2.2) is the filtered fiber bundles across all the merged ROI pairs for each scan $\mathcal{F}^{(2)}_{it} = \bigcup_{v,v'=1}^{\widetilde{V}} \mathcal{S}^{(2)}_{v,v',i,t}$, where $\mathcal{S}^{(2)}_{v,v',i,t}$ is the filtered tractograph of fibers between $R^{(1)}_v$ and $R^{(1)}_{v'}$.

**Fiber centroid extraction via clustering**

In (**S2.3**), we cluster fiber curves within each ROI pair across subjects (**Supplementary Figure S16C**). That is, we utilize the shape features of fiber curves to integrate and classify the tractograph patterns across subjects and build a common initial brain fiber atlas to capture the major characteristics of the structural connectome in the large population. One major output of this step is the clustering results $(\mathcal{C}^k_{v,v'})_{k,v,v'}$ of $\bigcup_{i=1}^n \mathcal{S}^{(2)}_{v,v',i,t_{i,1}} = \bigcup_{k=1}^{C_{v,v'}} \mathcal{C}^k_{v,v'}$, where $\mathcal{C}^k_{v,v'}$ is the $k$-th cluster that connects the ROI pair $(R_v, R_{v'})$, $t_{i,1}$ is the baseline observation time of subject $i$, and $C_{v,v'}$ is its total number of clusters between this ROI pair. If the number of fiber tracks connecting $R_v$ and $R_{v'}$ is large, then it is important to identify these pathways across subjects and scans and refine the parcellation such that each ROI pair only contains the major fiber pathways. We use the Tract Dictionary Learning (TractDL) as described in (30) to cluster the fibers. Specifically, each fiber is mapped from the native point space to a Hilbert space and then represented by the linear combination of a series of the cosine basis functions (31). This representation only depends on the degrees of the cosine basis functions and robust to the number of sampling points along the fiber. Tract Dictionary Learning (32) is further performed based on the cosine coefficients to learn the dictionary for each bundle. The $\mathbb{L}_1$ penalty is added to enforce each dictionary has sparse coefficients. Finally, each fiber is classified to a fiber bundle by minimizing the distance to each bundle dictionary.

**Selection of number of clusters**

In $(\mathbf{S2.3})$, we choose the number of clusters $C_{v,v'}$ to be proportional to the average number of fibers between each ROI pair. Specifically, $C_{v,v'} = \lceil (m_{v,v'} - m_1)/(\max_{v_1,v_2} m_{v_1,v_2} - m_1) \times (N_0 - 1) \rceil + 1$, where $m_{v,v'} = \sum_{i=1}^{n} N_{(v,v'),i,t_{i,1}}/n$ is the average fiber number between the ROIs $R_v$ and $R_{v'}$; $m_1 = \max\{\min_{v_1,v_2} m_{v_1,v_2}, m_0\}$; $m_0$ and $N_0$ serve as predefined thresholds, with $m_0$ representing the maximum allowable fiber count within a single cluster, and $N_0$ denoting the upper limit for the number of clusters across all pairs of regions of interest (ROIs); for any real number x, $\lceil x \rceil$ denotes the smallest integer that is no less than x. For example, in the HCP-Y data, the maximum fiber counts across ROI pairs averaged across subjects $m_1 = 7752$ fibers. For the number of clusters, we chose the maximum clustering number to be $N_0 = 99$ clusters, and the maximum allowable fiber counts in a single cluster to be $m_0 = m_1/100$. Clustering results for ROI pairs within the right somatomotor cortex (the R-3b and R-1 merged and R-OP4), prefrontal cortex (R-p10p and R-p47r), prefrontal cortex (R-8BL) and right putamen, left superior parietal and somatosensory cortices (L-2 and L-3b), brainstem and right putamen, left and right primary motor cortices (L-4 and R-4), left and right prefrontal cortices (L-9m and R-9m) and left and right somatosensory cortices (L-3b and R-5m) are depicted in eight subplots of **Supplementary Figure S16C**. They are presented in a clockwise sequence beginning from the top, respectively. Colors are used to visually differentiate major fiber clusters.

**Centroid fibers for atlas construction**

In $(\mathbf{S2.3})$, after we cluster fibers between all ROI pairs, we extract the central fiber from each cluster: we define the central fiber of a cluster $\mathcal{C}^k_{v,v'}$ as the fiber that has the smallest median MDF distance from all the other fibers inside the cluster. In particular, the central fiber of each cluster is defined as $S^k_{v,v'} = \arg\min_{s_{v,v'} \in \mathcal{C}^k_{v,v'}} \text{Median}_{u_{v,v'} \neq s_{v,v'}, u_{v,v'} \in \mathcal{C}^k_{v,v'}} [\text{MDF}(s_{v,v'}, u_{v,v'})]$. The final output of this step is the raw fiber skeleton atlas $\bigcup_{v,v'=1}^{\widetilde{V}} \bigcup_{k=1}^{C_{v,v'}} \left\{ \left( R_v^{(1)}, R_{v'}^{(1)} \right) \oplus S^k_{(v,v')} \right\}$.

**Post-clustering QC**

In $(\mathbf{S2.4})$, we performed post-clustering quality control based on cluster size and reproducibility. Clusters containing very few streamlines were discarded **Supplementary Figure S16D**. For the remaining clusters $\mathcal{C}^k_{v,v'}$ across all ROI pairs, the corresponding centroid fibers $S^k_{(v,v')}$ were selected to form a raw fiber atlas. Although this step is part of Stage 2, its outputs were further evaluated in the following stage $(\mathbf{S3.1})$, where subject-level FA profiles were projected onto the raw atlas and used to evaluate the ICC of FA profiles on centroid fibers based on 2,890 UKB test-retest subjects; low-quality clusters with ICC of FA less than $0.4$ and their centroid fibers were then excluded from the atlas. Specifically, fibers were removed if the projected FA profiles along the fiber trajectory showed low reliability (23). Applying these criteria yielded the final refined fiber clusters $(\widetilde{\mathcal{C}}^k_{v,v'})_{v,v',k}$ and their corresponding centroid fibers $\widetilde{S}^k_{(v,v')}$, for $v, v' = 1, \ldots, \widetilde{V}$ and $k = 1, \ldots, \widetilde{C}_{v,v'}$, where $\widetilde{C}_{v,v'}$ denotes the number of clusters connecting $R_v$ and $R_{v'}$ after post-clustering quality control. The output of this step was the fiber skeleton atlas (Fig. 1B and Supplementary Fig. S3), also referred to as the parcellation-based tractographic skeleton (CBSS), defined as

$$\text{CBSS} = \bigcup_{v,v'=1}^{\widetilde{V}} \left\{ \left( R_v^{(1)}, R_{v'}^{(1)} \right) \oplus \bigcup_{k=1}^{\widetilde{C}_{v,v'}} \widetilde{S}_{(v,v')}^{(k)} \right\}.$$

The total number of fibers in the final atlas was $M = \sum_{v,v'=1}^{\widetilde{V}} \widetilde{C}_{v,v'}$. The raw fiber atlas initially contained 6,090 fibers, which was reduced to $M = 5,723$ fibers after post-clustering QC.

In $(\mathbf{S2}.4)$ we perfrom QC after clustering, based on cluster size and reproducibility. Clusters containing an extremely small number of fibers are discarded **Supplementary Figure S16D**. Then central fibers $S_{(v,v')}^k$ from the remaining cluster $\mathcal{C}_{v,v'}^k$ across all ROI pairs are selected to form the raw fiber atlas. The FA values from each subject are projected to the raw fiber atlas, according to steps in the next section (S3.1). Fibers are excluded if their reliability (23) of the projected FAs along fiber points are too small. Applying these criteria will result in the final fiber clusters $(\widetilde{\mathcal{C}}_{v,v'}^k)_{v,v',k}$ and their corresponding centroid fibers $\widetilde{S}_{(v,v')}^k$, for $v, v' = 1, \cdots, \widetilde{V}, k = 1, \cdots, \widetilde{C}_{v,v'}$ where $\widetilde{C}_{v,v'}$ is the number of clusters connecting $R_v$ and $R_{v'}$ after the post-clustering QC. The final output of this step is the fiber skeleton atlas (**Figures 1B**, and **Supplementary Figure S3**), or the parcellation-based tractographic skeleton (CBSS), which is given by CBSS $= \cup_{v,v'=1}^{\widetilde{V}} \left\{ \left( R_v^{(1)}, R_{v'}^{(1)} \right) \oplus \cup_{k=1}^{\widetilde{C}_{v,v'}} \widetilde{S}_{(v,v')}^{(k)} \right\}$. The total fiber number is $M = \sum_{v,v'=1}^{\widetilde{V}} \widetilde{C}_{v,v'}$. The 6090 fibers included in the raw fiber skeleton atlas are reduced to $M = 5723$ fibers in the final refined atlas after post-clustering QC. Next we will list detailed substeps in stage three: (S3.1) Fiber projection for downstream cohorts; (S3.2) elastic registration; and (S3.3) Low-dimensional representations of tractographic data.

**Fiber projection for downstream cohorts**

The input of (S3.1) is the nonlinearly registered spatial statistics following the TBSS pipeline (33). Remember from the TBSS pipeline, the FA images from all subjects are aligned, undergoing first an affine transformation and then a nonlinear registration into the $1 \times 1 \times 1$ mm$^3$ MNI152 space. All further processing is conducted within this space and resolution for ease of interpretation and visualization. The spatial statistics projected can include the DTI maps FA, MD, MO, axial diffusivity (AD), radial diffusivity (RD). First we will project those voxel-based FA maps onto our established atlas as an alignment-invariant tract representation, by filling the fiber atlas skeleton with FA values from the nearest relevant tract centre. Similar to (33), this is achieved, for each skeleton voxel, by searching perpendicular to the local fiber skeleton structure for the maximum value in the subject's FA image. Specifically, Skeletonization is achieved by initially determining the direction perpendicular to the local surface at every voxel within the image; non-maximum suppression is applied in this perpendicular direction; a search is conducted through the voxels in the direction perpendicular to the tract within the local $1 \times 1 \times 1$ voxel neighbourhood, and the voxel exhibiting the highest FA value is identified as the tract's central point (**Supplementary Figure S16E**); the coordinates of the searched voxel with the maximum FA will be recorded, and other spatial statistics will be extracted at these coordinates. Here we use $1 \times 1 \times 1$ voxel neighbourhood in the fiber skeletonization, which is different from the $3 \times 3 \times 3$ voxel neighbourhood of the TBSS pipeline, to capture fine details of fibers and reduce overlap between fibers as fiber atlas is spatially dense. In addition, for each voxel on the fibers, we do trilinear interpolation of FA to resample its neighborhood to be of the resolution $0.1 \times 0.1 \times 0.1$ before we make the FA projection. The final output of this step is $\widetilde{\mathcal{P}}_{it} = \left\{ Y_{it,(v,v'),k}^{(D)}(s) | k \leq \widetilde{C}_{v,v'}, v, v' \leq \widetilde{V}, s \in [0,1] \right\}$ consists of $M$ one-dimensional functions $Y_{it,(v,v'),k}^{(D)}(\cdot)$ which represents the diffusion profiles $(D)$

such as FA, projected onto the fiber tract $\widetilde{S}^{(k)}_{(v,v')}$. Compared to the FA map at individual tractography space $\mathcal{P}_{it}$, projecting the FA onto the common atlas facilitate fiber-level voxel-based population analysis.

**Elastic fiber registration.**

In (**S3.2**), we performed group-wise elastic registration of fiber-wise diffusion profiles within each ROI pair across subjects. Existing streamline-registration methods (34–40) can suffer from limitations such as pinching or unstable pointwise correspondence; see (41) for discussion. To address this, we used an elastic functional registration framework based on the square-root velocity (SRV) representation, which provides a reparameterization-invariant metric for curve alignment under the elastic geometry of functions (41).

Let

$$Y^{\mathcal{D}}_{i,t} = \left\{ Y^{(D)}_{i,t,(v,v'),j}(s) : v, v' \leq \widetilde{V},\ j \leq \widetilde{C}_{v,v'},\ D \in \mathcal{D},\ s \in [0,1] \right\}$$

denote the raw tract-based diffusion features for subject $i$ at time $t$, where $Y^{(D)}_{i,t,(v,v'),j}(s)$ is diffusion parameter $D$ measured at location $s$ along centroid fiber $S^{(j)}_{(v,v')}$. Although diffusion maps were already nonlinearly registered to MNI space in (**S1.1**), residual fiber-level misalignment may remain because homologous diffusion patterns can be shifted or stretched unequally across subjects. We therefore aligned the fiber-wise FA profiles across subjects before downstream feature extraction.

For each centroid fiber $S^{(j)}_{(v,v')}$, we considered the FA profile $Y^{FA}_{i,t,(v,v'),j}(s)$ and its SRV representation

$$q_{Y^{FA}_{i,t,(v,v'),j}}(s) = \frac{\dot{Y}^{FA}_{i,t,(v,v'),j}(s)}{\sqrt{|\dot{Y}^{FA}_{i,t,(v,v'),j}(s)|}},$$

where $\dot{Y}$ denotes the derivative with respect to $s$. Let $\bar{Y}^{FA}_{(v,v'),j}(s)$ denote the cross-subject Karcher mean profile for fiber $S^{(j)}_{(v,v')}$. For subject $i$, we estimated a smooth warping function $\gamma_{i,(v,v'),j} \in \Gamma$, where $\Gamma$ is the set of boundary-preserving monotone diffeomorphisms of $[0,1]$, by minimizing the penalized elastic objective

$$\gamma_{i,t,(v,v'),j} = \arg\min_{\gamma \in \Gamma} \left[ \int_0^1 \left\{ q_{Y^{FA}_{i,t,(v,v'),j}}(\gamma(s))\sqrt{\dot{\gamma}(s)} - q_{\bar{Y}^{FA}_{(v,v'),j}}(s) \right\}^2 ds + \lambda\, l_{S^{(j)}_{(v,v')}} \int_0^1 \{\gamma(s) - s\}^2 ds \right].$$

The first term measures the SRV-based elastic discrepancy between the warped subject-specific profile and the group mean, whereas the second term penalizes excessive deviation from the identity map. Here $l_{S^{(j)}_{(v,v')}}$ denotes the streamline length of $S^{(j)}_{(v,v')}$, and $\lambda$ is a tuning parameter controlling the amount of regularization. This formulation allows nonlinear phase variation to be removed while discouraging biologically implausible warping. In practice, optimization was carried out iteratively within each ROI pair by alternating between estimation of the group mean profile and subject-specific warping functions until convergence.

The aligned diffusion profiles were then defined as

$$(Y \circ \gamma)^{\mathcal{D}}_{i,t} = \left\{ Y^{(D)}_{i,t,(v,v'),j} \circ \gamma_{i,t,(v,v'),j} \right\}_{v,v',j,D \in \mathcal{D}},$$

where $\circ$ denotes function composition. **Supplementary Figure S16F** illustrates the effect of elastic alignment for a representative tract cluster. Based on $(Y \circ \gamma)^{\mathcal{D}}_{i}$, we derived multiscale edge features. We extracted summary measures of aligned diffusion profiles, such as the mean, to obtain

$$Y_{i,t}^{W} = \left(Y_{i,t,(v,v')}^{W}\right)$$

to characterize the fiber-level mean FA-based structural connectivity for the pathway between $R_v$ and $R_{v'}$ for subject $i$.

**Low-dimensional representations of tractographic data**

The (S3.3) consists of both basis learning and efficient representation based on a matrix of basis functions across all pairs of ROIs across time. After alignment, all fiber tracts connecting a specific ROI pair across subjects and time share many shape similarities. Without loss of generality, we focus on the $k-$th representative fiber (cluster centroid) for an ROI pair, $(\widetilde{R}_v, \widetilde{R}_{v'})$. We perform functional principal component analysis (fPCA) on the FA profiles $Y_{it,(v,v'),k}^{\mathcal{D}}(s) = Y_{it,(v,v'),k}^{(FA)}(s)$ of each aligned fiber tract across subjects, in order to learn a set of basis functions denoted as $\mathcal{L}_{(v,v',k)} = \left\{\phi_{(v,v',k),m} : m = 1, \ldots, M\right\}$ , where $M$ is the number of fPCA basis and $\phi_{(x,y,z,t),m}$ are the fPCA basis functions. Each fiber curve connecting $\left(\widetilde{R}_v, \widetilde{R}_{v'}\right)$ at time $t$ can be sparsely represented as a linear combination of the basis functions in $\mathcal{L}_{(v,v')}$. We then projected the FA profiles $Y_{it,(v,v'),k}(s)$ onto the PC bases to obtain PC scores (embeddings) extracted network-level hiearchical PCs.

In $(\mathbf{S3.3})$, we performed basis learning and efficient representation of aligned fiber-wise FA profiles. Without loss of generality, we focus on the $k$-th representative fiber (cluster centroid) for ROI pair $(\widetilde{R}_v, \widetilde{R}_{v'})$. For the aligned FA profiles $Y_{it,(v,v'),k}(s)$, we performed functional principal component analysis (fPCA) across subjects to learn a fiber-specific basis set

$$\mathcal{L}_{(v,v',k)} = \{\phi_{(v,v',k),m}(s) : m = 1, \ldots, M_{(v,v',k)}\},$$

where $\phi_{(v,v',k),m}(s)$ denotes the $m$-th eigenbasis function and $M_{(v,v',k)}$ is the retained number of fiber-level bases. The corresponding mean profile is denoted by $\mu_{(v,v',k)}(s)$. Each FA profile was then represented as

$$Y_{it,(v,v'),k}(s) = \mu_{(v,v',k)}(s) + \sum_{m=1}^{M_{(v,v',k)}} \xi_{it,(v,v'),k,m}\, \phi_{(v,v',k),m}(s) + \varepsilon_{it,(v,v'),k}(s),$$

where

$$\xi_{it,(v,v'),k,m} = \int \{Y_{it,(v,v'),k}(s) - \mu_{(v,v',k)}(s)\}\, \phi_{(v,v',k),m}(s)\, ds$$

is the $m$-th fiber-level PC score for subject $i$ at time $t$. Thus, each fiber tract was encoded by the fiber-level embedding vector

$$\boldsymbol{\xi}_{it,(v,v'),k} = (\xi_{it,(v,v'),k,1}, \ldots, \xi_{it,(v,v'),k,M_{(v,v',k)}})^{\top}.$$

To obtain network-level representations, let $\mathcal{N}_a$ and $\mathcal{N}_b$ denote two functional networks, and let

$$\mathcal{P}_{ab} = \{(v, v') : \widetilde{R}_v \in \mathcal{N}_a,\ \widetilde{R}_{v'} \in \mathcal{N}_b\}$$

be the set of ROI pairs belonging to network pair $(\mathcal{N}_a, \mathcal{N}_b)$. We concatenated all fiber-level PC scores from ROI pairs in $\mathcal{P}_{ab}$ to form a network-pair feature vector

$$\mathbf{z}_{it,ab} = \mathrm{vec}\big\{\boldsymbol{\xi}_{it,(v,v'),k} : (v, v') \in \mathcal{P}_{ab},\ k = 1, \ldots, \widetilde{C}_{v,v'}\big\},$$

where $\widetilde{C}_{v,v'}$ is the number of retained centroid fibers for ROI pair $(v, v')$. We then performed a second-stage PCA on $\mathbf{z}_{it,ab}$ across subjects to obtain a set of hierarchical basis vectors

$$\mathcal{H}_{ab} = \{\psi_{ab,\ell} : \ell = 1, \ldots, L_{ab}\},$$

where $\psi_{ab,\ell}$ is the $\ell$-th hierarchical eigenbasis for network pair $(\mathcal{N}_a, \mathcal{N}_b)$ and $L_{ab}$ is the retained number of hierarchical components. The resulting hierarchical PC scores were

$$\eta_{it,ab,\ell} = \mathbf{z}_{it,ab}^{\top}\psi_{ab,\ell}, \qquad \ell = 1, \ldots, L_{ab}.$$

Therefore, each subject and time point was represented at the network-pair level by the hierarchical embedding

$$\boldsymbol{\eta}_{it,ab} = (\eta_{it,ab,1}, \ldots, \eta_{it,ab,L_{ab}})^{\top}.$$

These hierarchical PC scores were used as the network-level features in downstream prediction analyses. The output of this stage therefore includes a graph of fiber-level basis functions and scores $\mathrm{GPCA}_{Fib}$, together with the network-level hierarchical bases and scores $\mathrm{GPCA}_{Net}$, with

$$\mathrm{GPCA}_{Fib} = \bigcup_{v,v'=1}^{\widetilde{V}} \left\{ \left(\widetilde{R}_v, \widetilde{R}_{v'}\right) \oplus \mathcal{L}_{(v,v')} \oplus \bigcup_{k=1}^{\widetilde{C}_{v,v'}} \left(\boldsymbol{\xi}_{it,(v,v'),k}\right)_{i,t} \right\},$$

and

$$\mathrm{GPCA}_{Net} = \bigcup_{a,b} \left\{ (\mathcal{N}_a, \mathcal{N}_b) \oplus \mathcal{H}_{ab} \oplus \left(\boldsymbol{\eta}_{it,ab}\right)_{i,t} \right\},$$

where $\mathcal{L}_{(v,v')} = \bigcup_{k=1}^{\widetilde{C}_{v,v'}} \mathcal{L}_{(v,v',k)}$ is the set of fiber-level fPCA bases for ROI pair $(\widetilde{R}_v, \widetilde{R}_{v'})$.

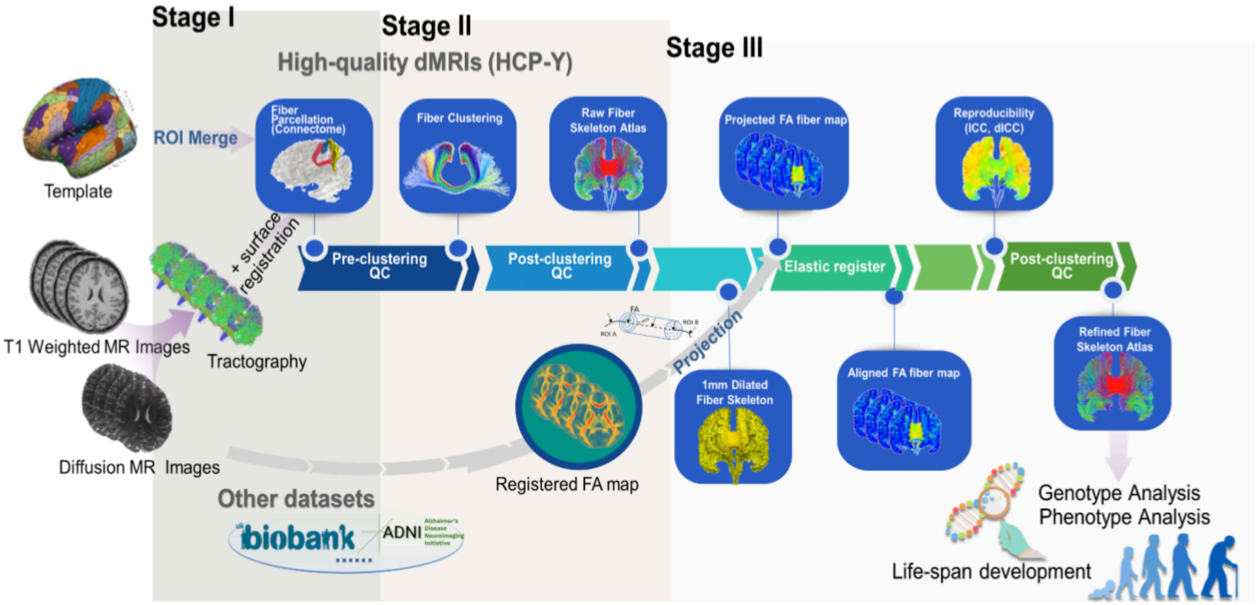


**Supplementary Figure S1. The overall dMRI data processing pipeline.**

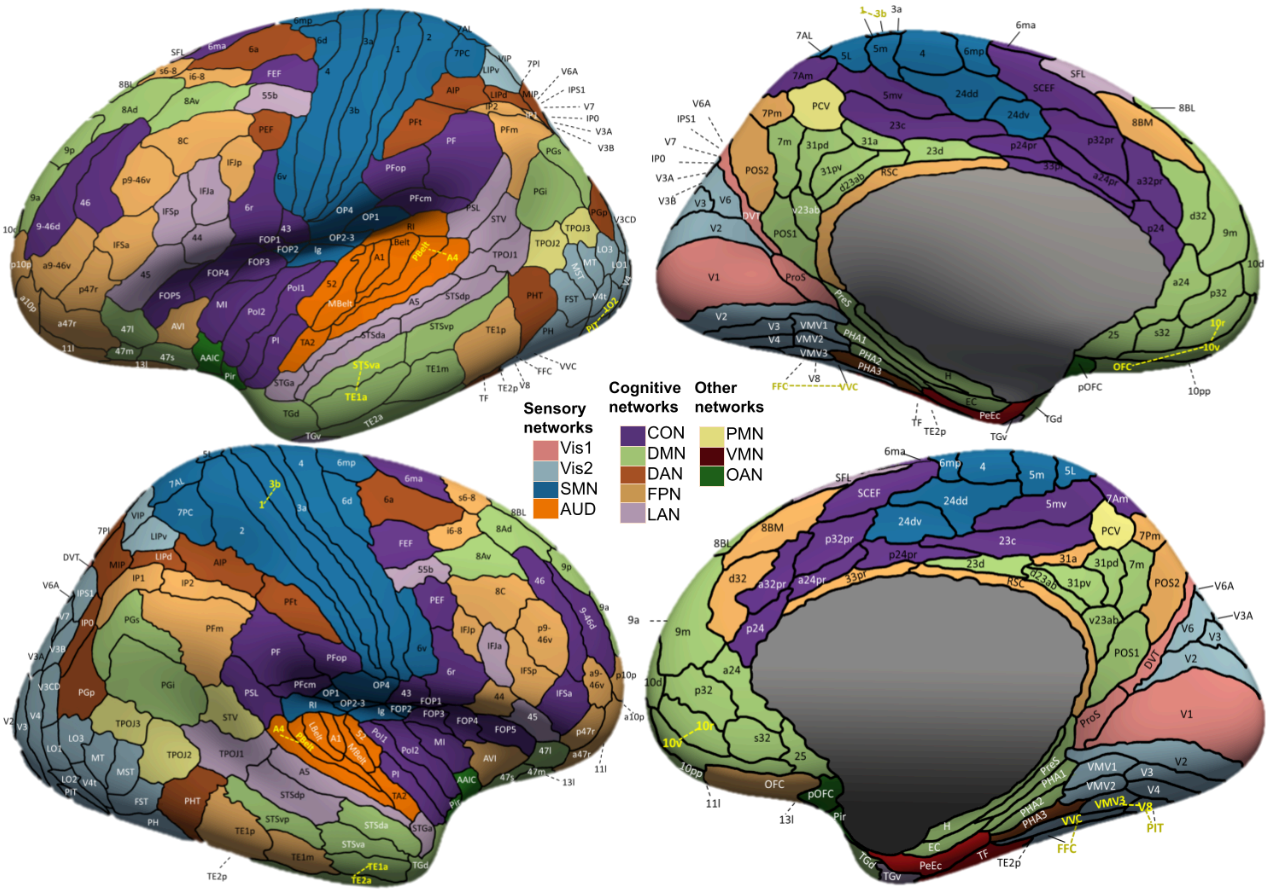


**Supplementary Figure S2. HCP-MMP cortical parcellation with merged ROIs highlighted.** Twenty-six parcels highlighted in yellow and brown were merged into 12 larger regions; dotted yellow/brown lines indicate the merged groups. ROI, region of interest. Merged ROIs comprised the following groups: secondary visual network, (L_LO2, L_PIT), (L_FFC, L_VVC), (R_V8, R_VMV3, R_PIT), (R_FFC,R_VVC); somatomotor network, (L_3b, L_1), (R_3b,R_1); default mode network, (L_10r, L_10v, L_OFC), (L_TE1a,L_STSva), (R_10r,R_10v), (R_TE1a, R_TE2a); and auditory network: (L_PBelt, L_A4), (R_PBelt, R_A4). The merging criteria were as follows: regions had to be spatially adjacent, belong to the same functional network, exhibit highly similar fiber-count profiles with other regions (correlation > 0.7), and be mergeable without significantly reducing test-retest reliability of fiber count via 43 pairs of HCP-Y test-retest scans.

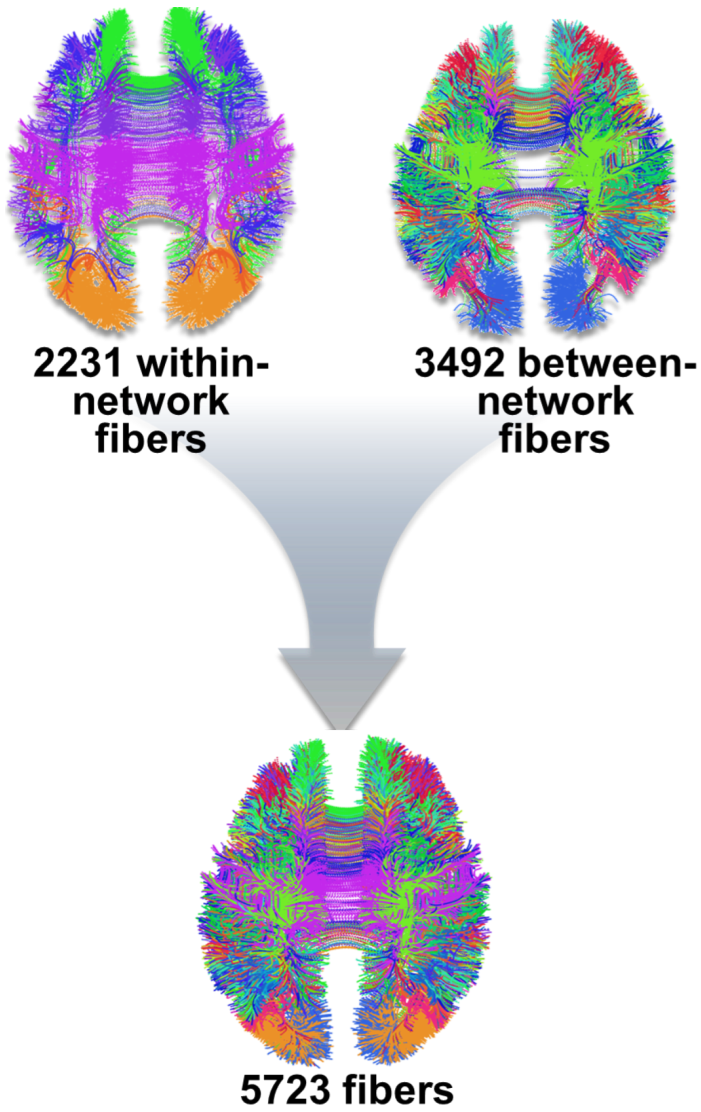


**Supplementary Figure S3. Between-network and within-network representative fibers in the BFSC atlas.** Colors indicate different network pairs.

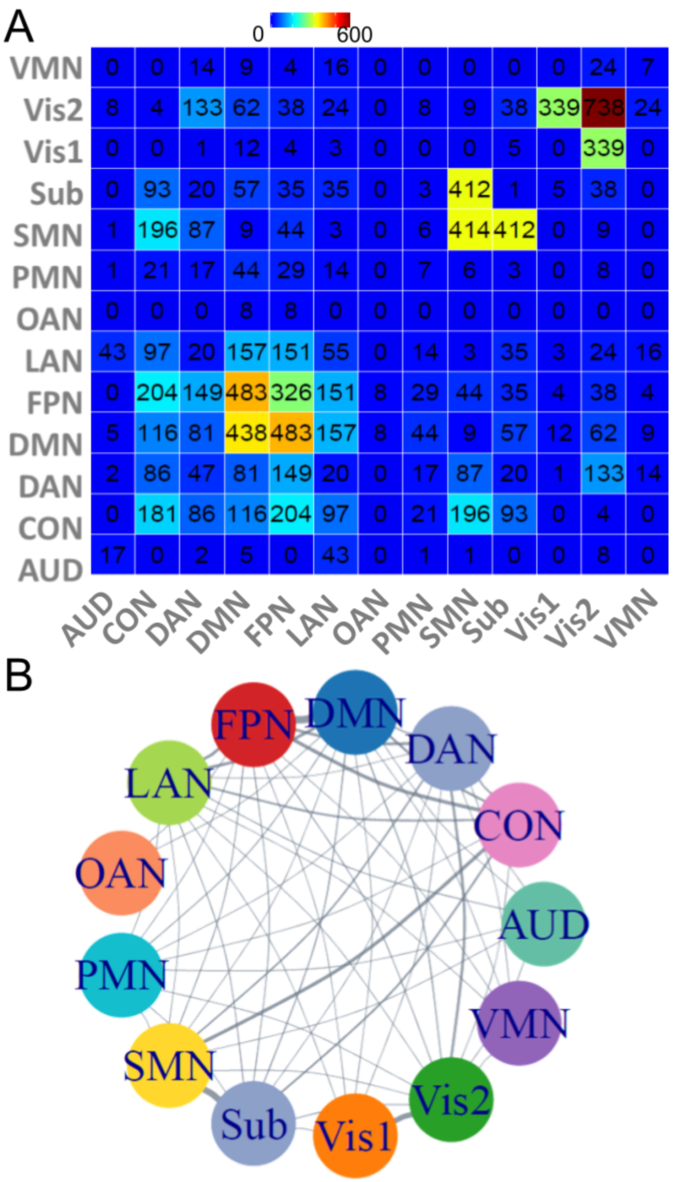


**Supplementary Figure S4. Fiber-count distribution of the BFSC atlas across network pairs.** **A.** Fiber-count distribution of the BFSC atlas across network pairs. **B.** Network graph showing the same pairwise counts, with edge width scaled by fiber count.

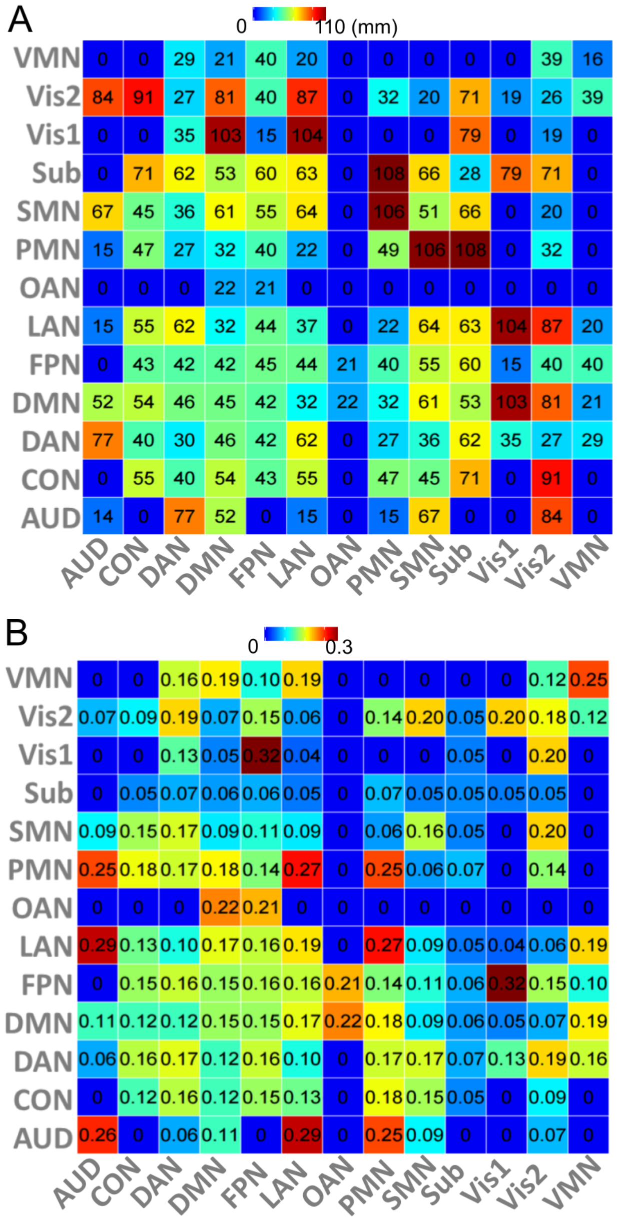


**Supplementary Figure S5. The mean fiber length (A) and curvature (B) of the BFSC atlas across all the 13 network pairs.**

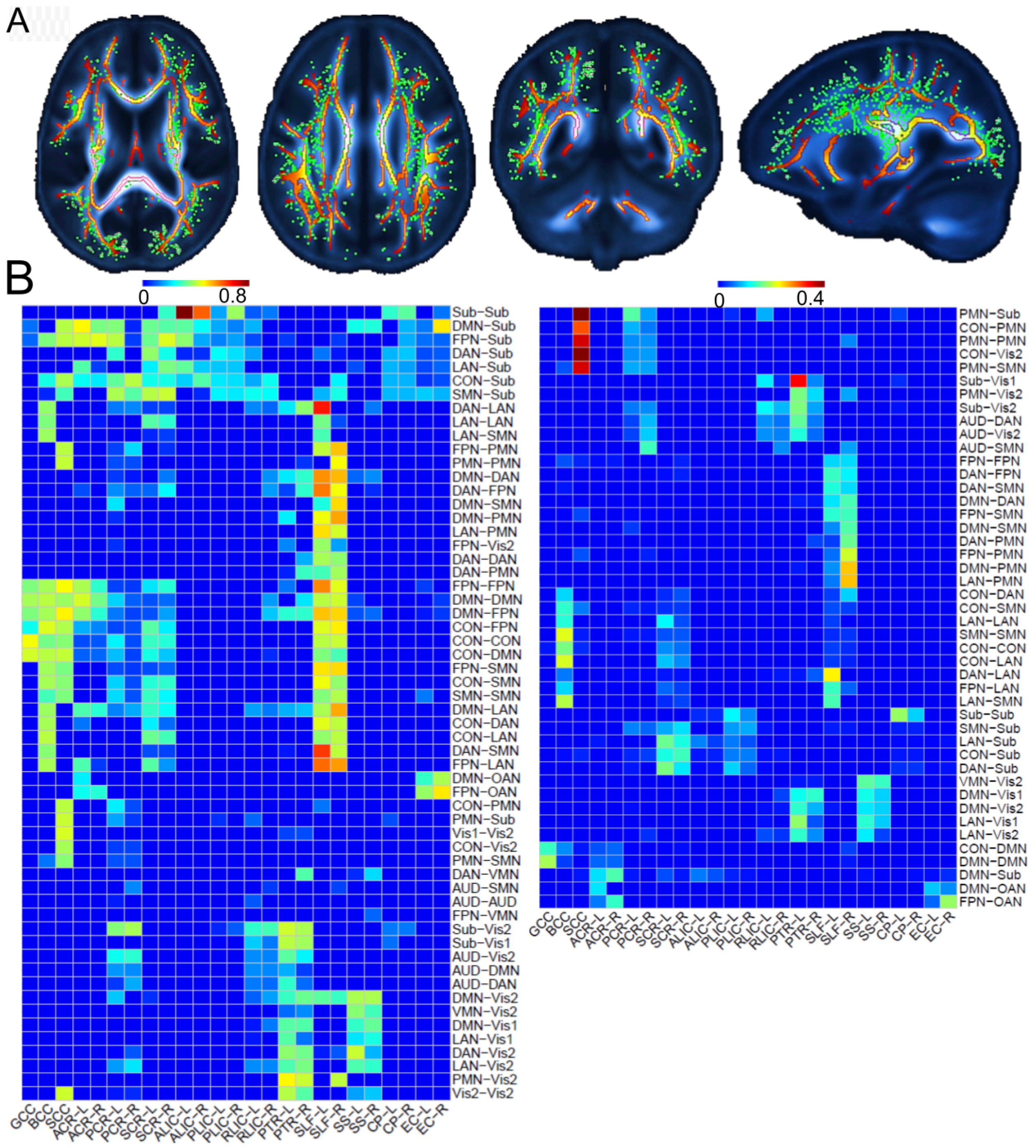


**Supplementary Figure S6. The overlap visualization of BFSC atlas (for CBSS) and the JHU atlas (for TBSS). A.** Fractional anisotropy (FA) template in MNI152 space with the TBSS skeleton overlaid as colored lines and BFSC atlas fiber points in the 2D slice shown as green dots. Representative axial, coronal, and sagittal views demonstrate the anatomical alignment. **B.** Overlap between BFSC network-pair fibers and JHU white-matter tracts. For each fiber, overlap with a JHU tract was quantified as the proportion of fiber length falling within the 1-mm dilation of that JHU tract, consistent with the 1-mm cylinder mapping used in CBSS. For each JHU tract and network pair, the left and right panels show the maximal and mean overlap percentages across all fibers linking that network pair, respectively.

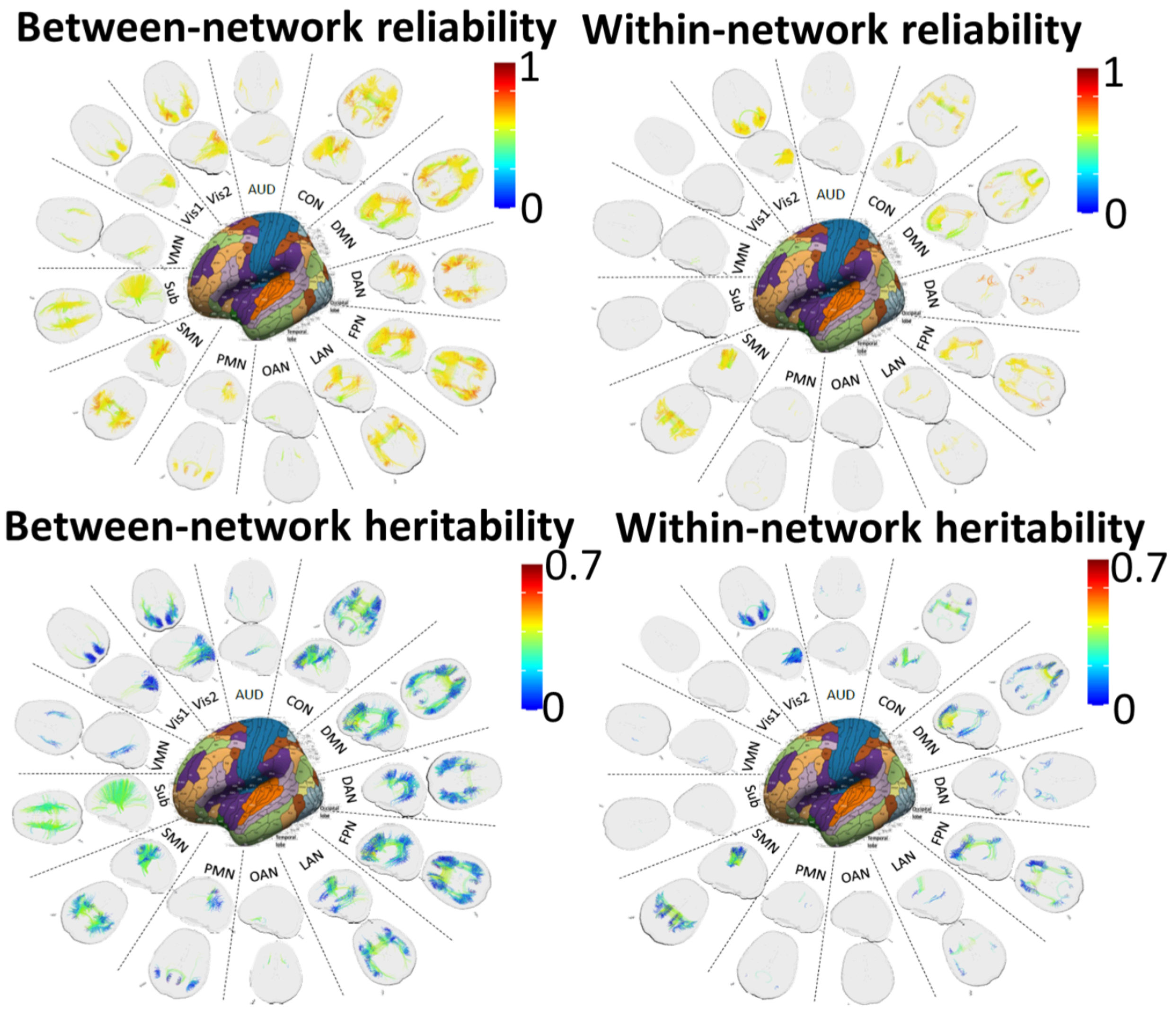


**Supplementary Figure S7. The visualization of fiber-level reliability and heritability. Upper,** The between-network and within-network fiber-level reliability. **Lower,** The between-network and within-network fiber-level heritability.

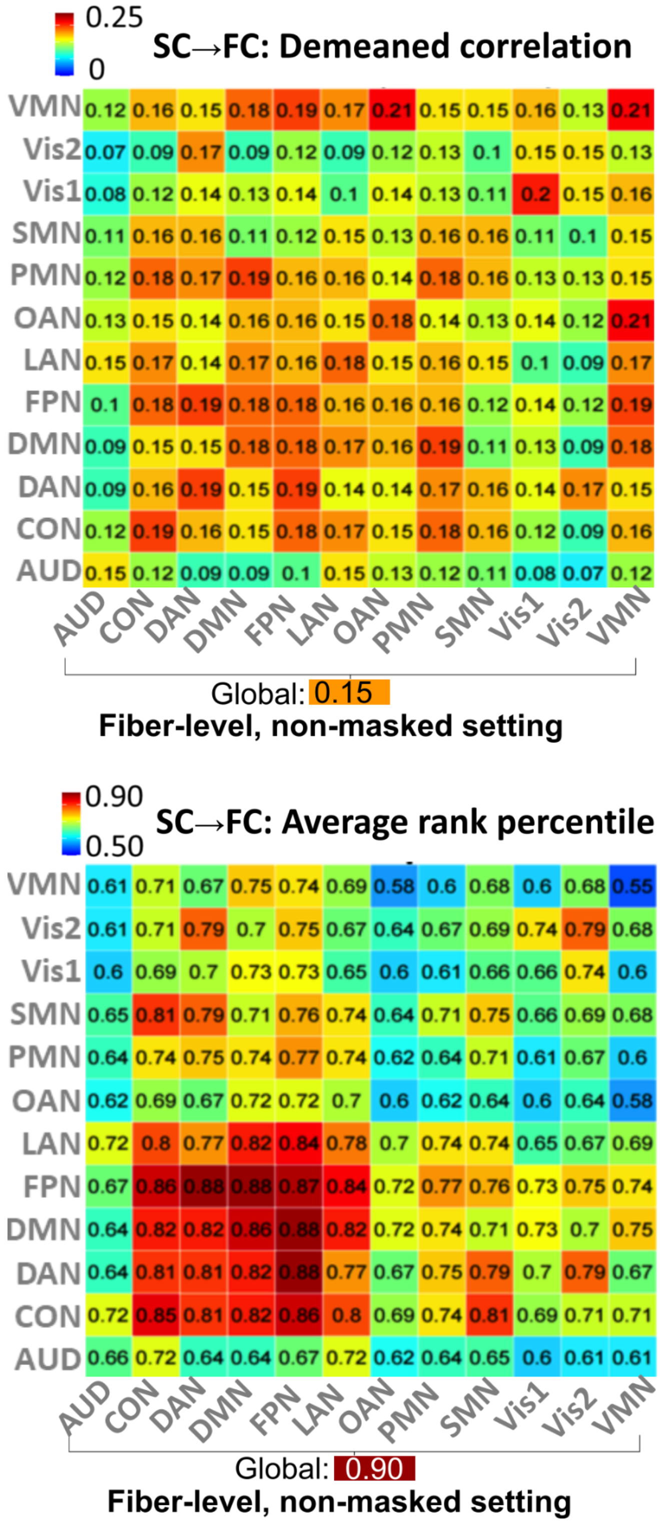


**Supplementary Figure S8. Functional connectivity prediction using fiber-level mean FA under the non-masked setting, in which all atlas fibers were used as predictors.** Top, de-meaned correlation; bottom, average rank percentile. FA, fractional anisotropy; SC, structural connectivity; FC, functional connectivity.

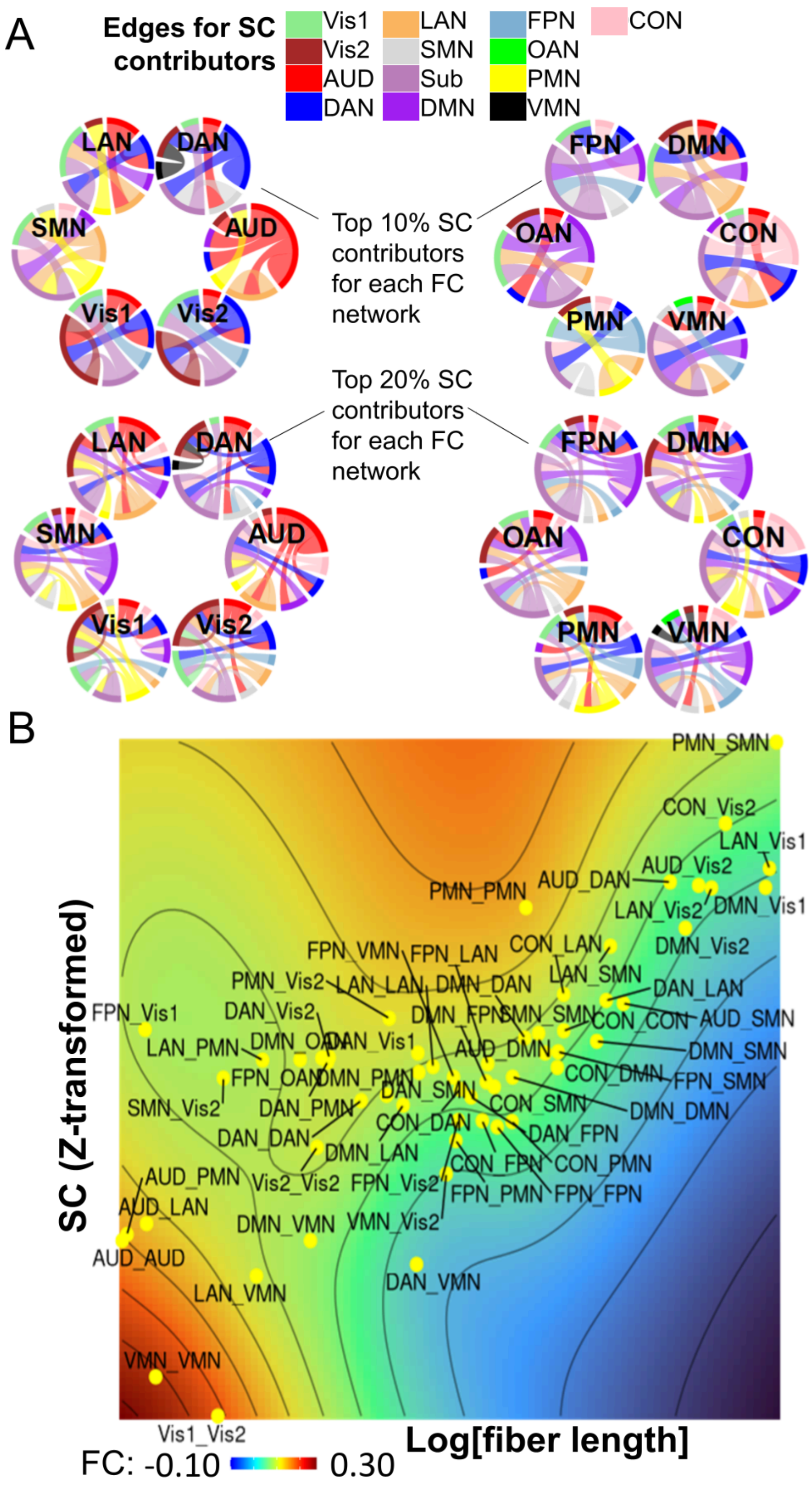


**Supplementary Figure S9. The visualization of structural-functional coupling. A.** The top 20% (middle circle) and top 10% signals (outer circle) for SC-FC coupling. **B.** The contour plot of all network-level FCs based on SCs (y-axis) and mean fiber length (x-axis).

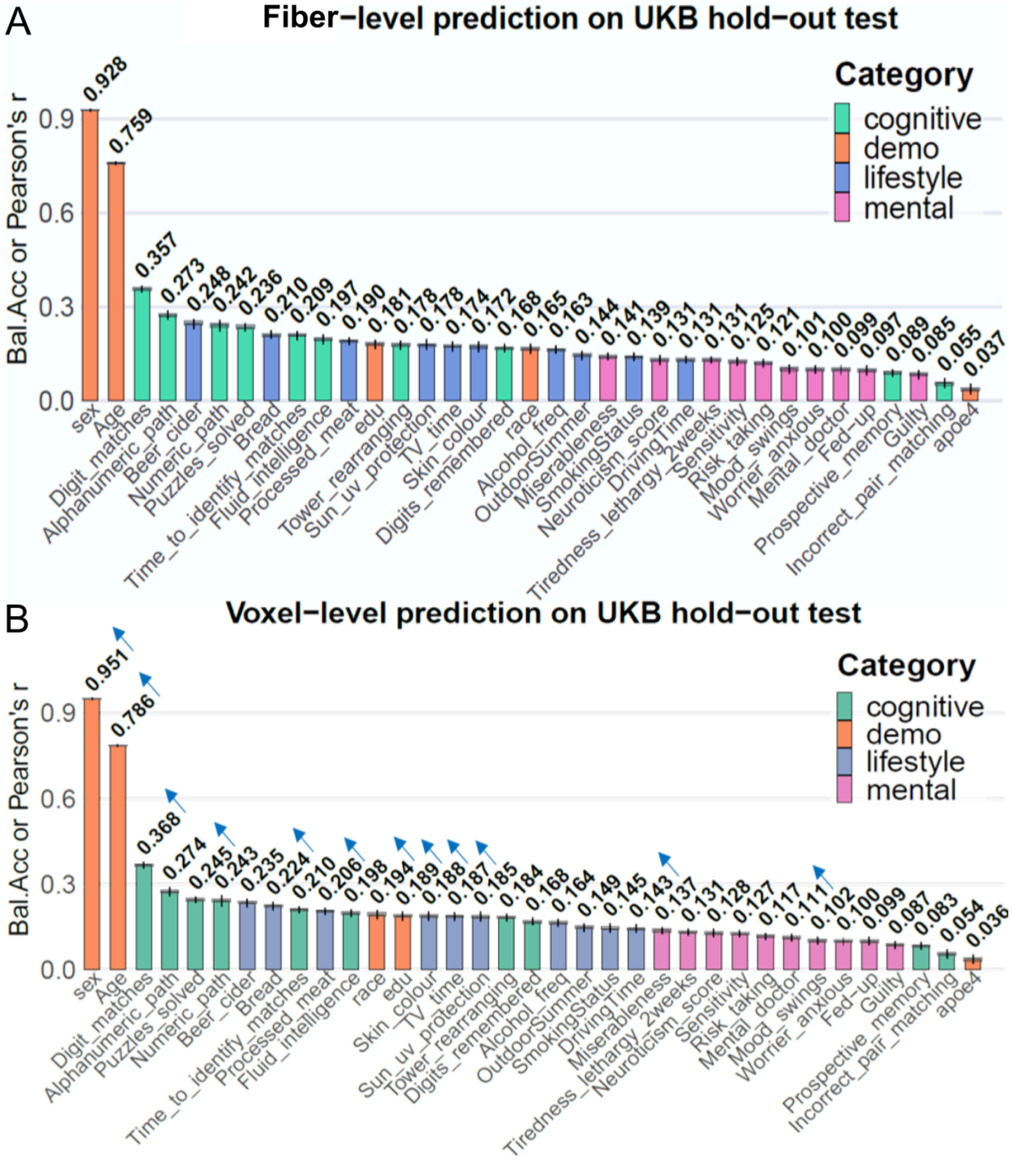


**Supplementary Figure S10. The performance for predictions of selected cognitive/behavioral traits based on fiber-level (A) and voxel-level (B) structural connectivities defined based on fractional anisotropy using the partial least square regression method.**

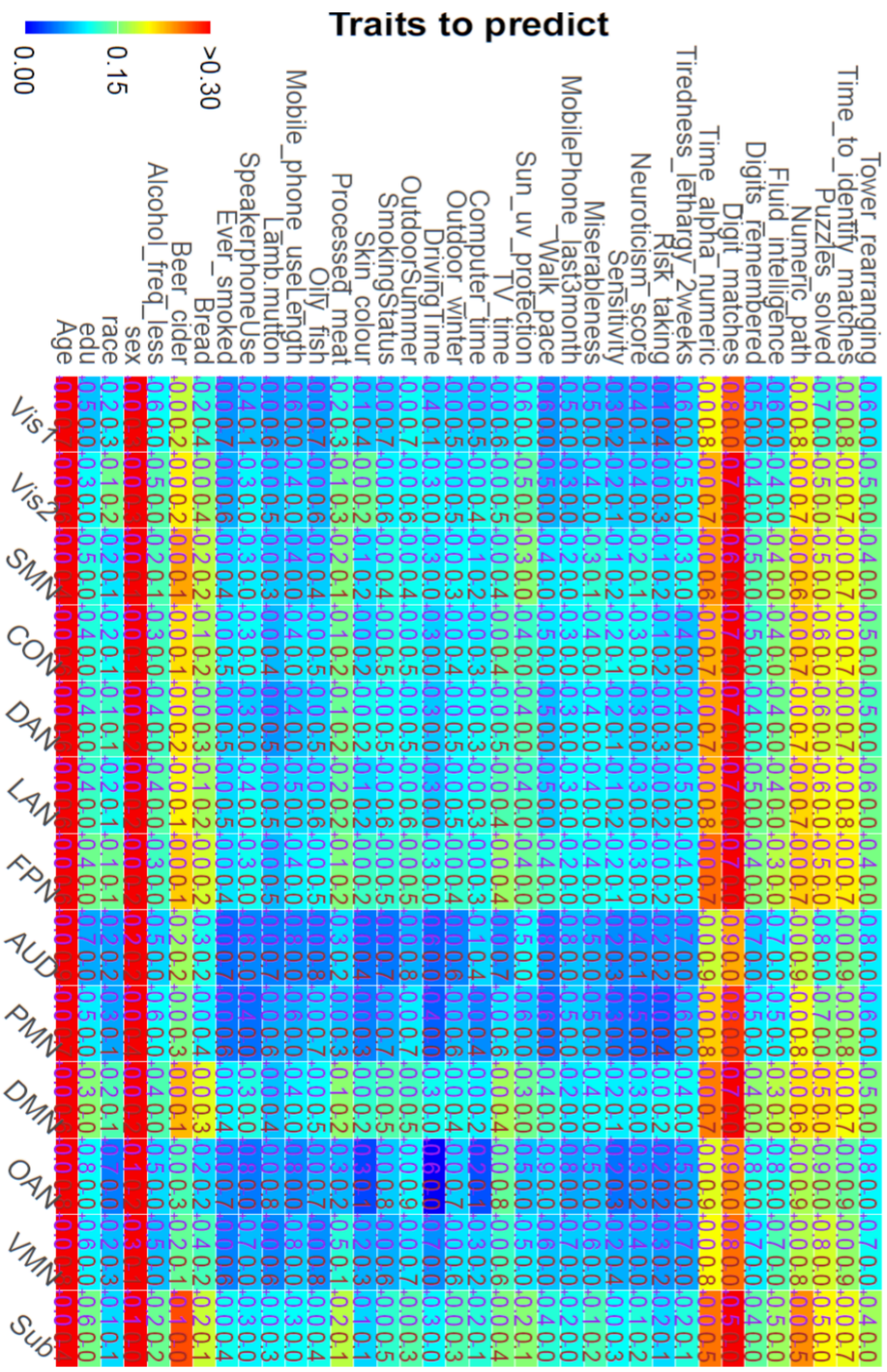


**Supplementary Figure S11. Signed SC contributions to cognitive and behavioral prediction using only pathways connected to each network.** For each cell, balanced accuracy or prediction correlation is shown separately by using SC features whose partial least-squares regression coefficients are positive-only (left) or negative-only (right), respectively. Values are set to 0 when predictions are non-significant or negatively correlated with the observed phenotypes. SC (structural connectivity) was defined using pathway-averaged FA (the fiber-level).

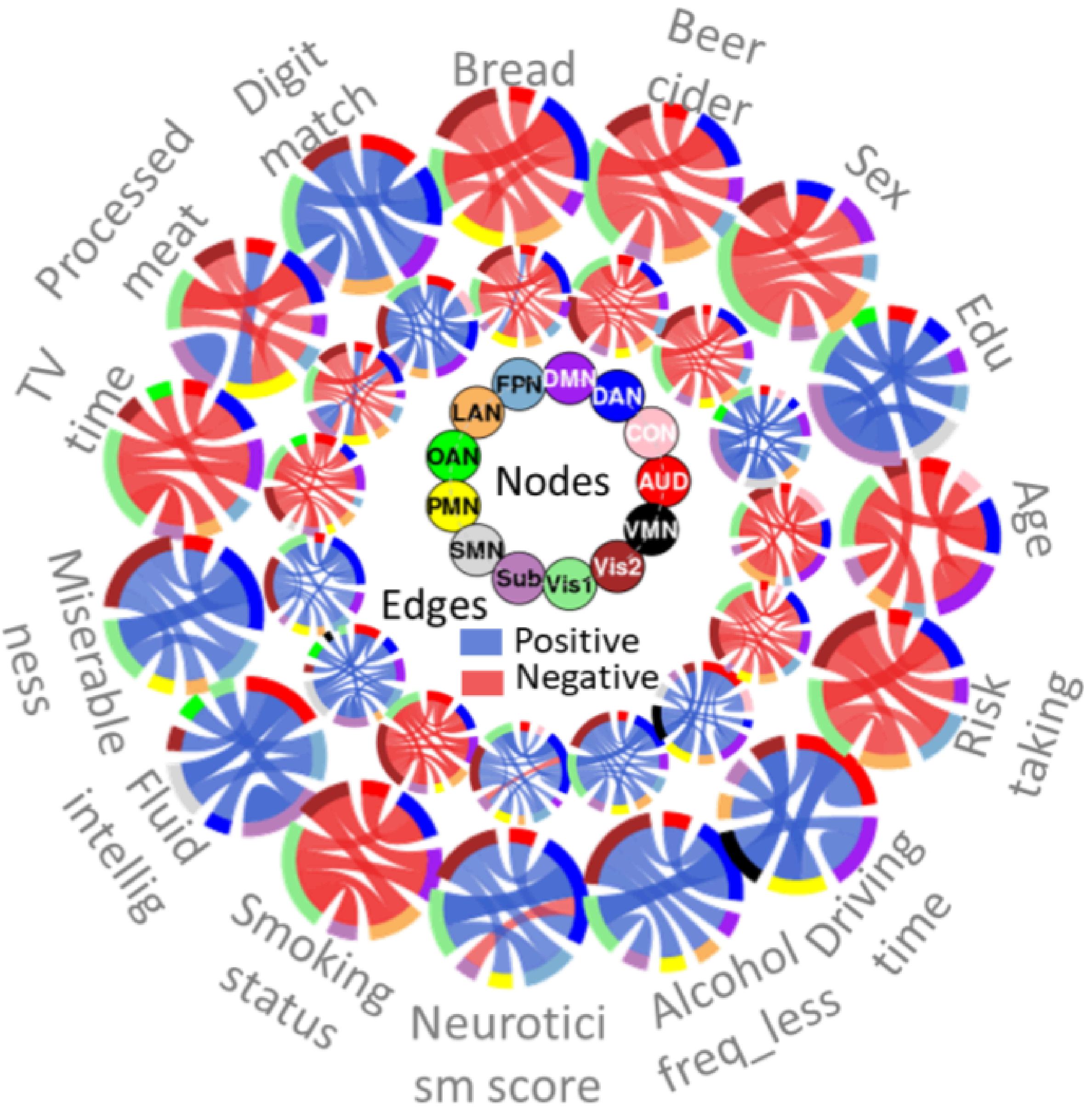


**Supplementary Figure S12. Top SC contributions.** The top 10% (outer ring) and top 20% (inner ring) of SC contributors for 15 selected traits. Red and blue indicate positive and negative associations, respectively. SC (structural connectivity) was defined using pathway-averaged FA (the fiber-level).

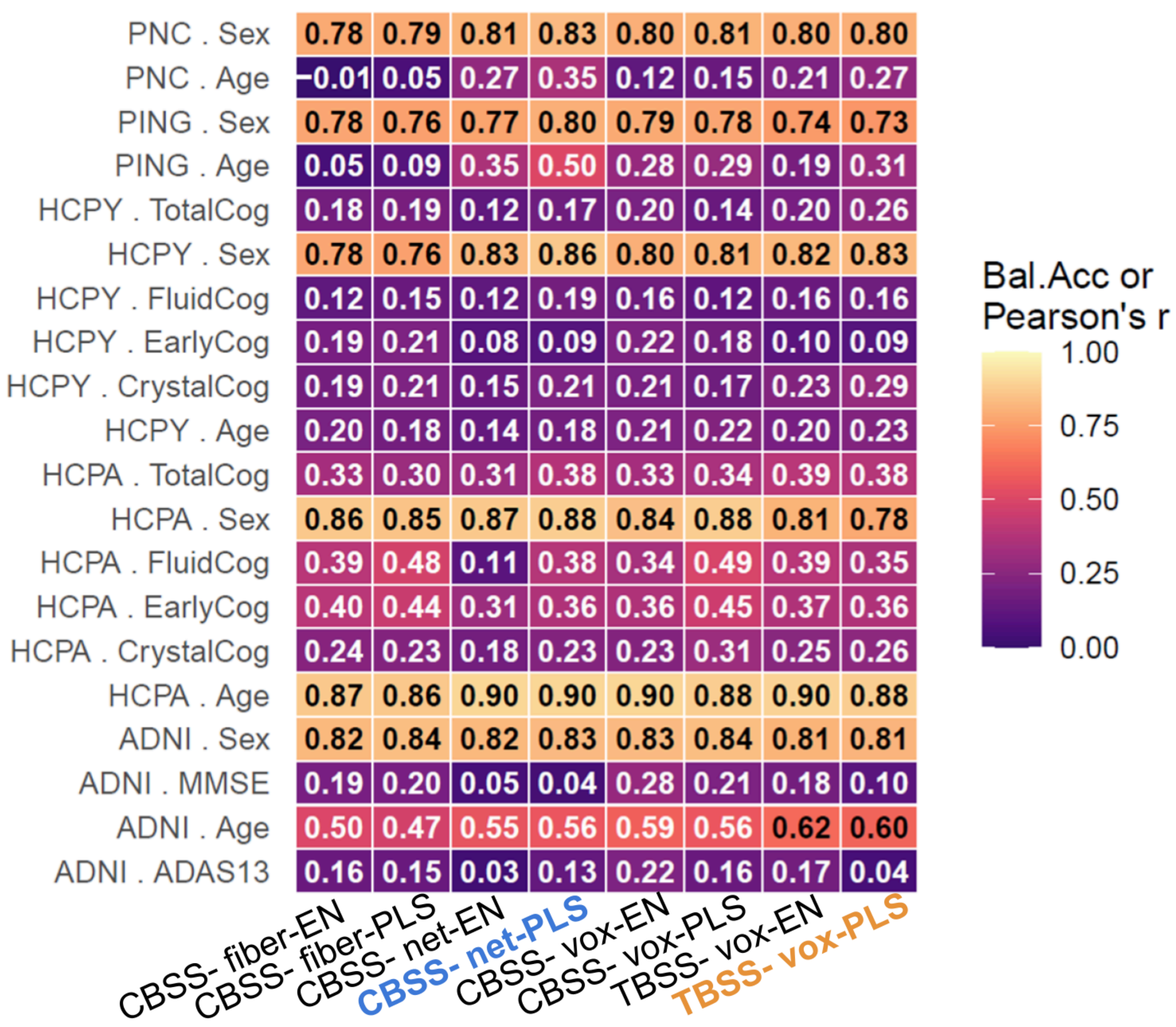


**Supplementary Figure S13. External validation across five lifespan cohorts by transferring UK Biobank-trained models for cognitive and behavioral prediction using fiber-, voxel- and network-level CBSS features and voxel-level TBSS features, evaluated with elastic net (EN) and partial least squares (PLS).** Models were transferred from the models trained by the UKB phase 1-2 data.

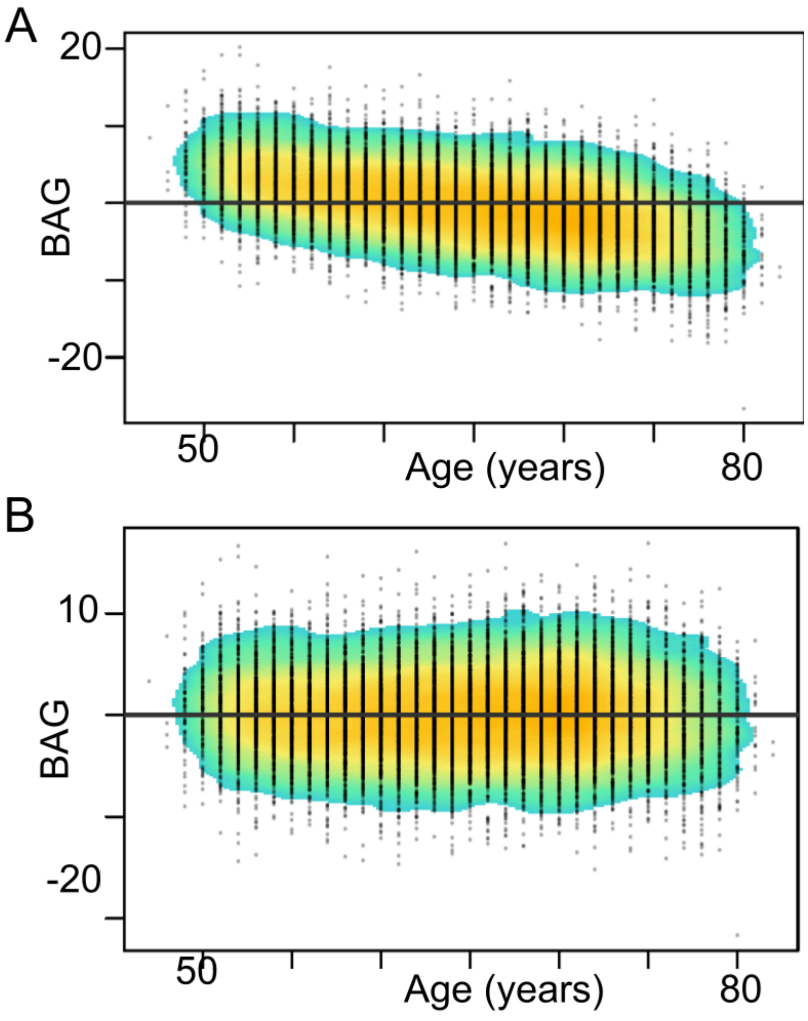


**Supplementary Figure S14. A comparison of scatterplots for brain age gap (BAG) with age, for before (A) and after (B) age correction.**

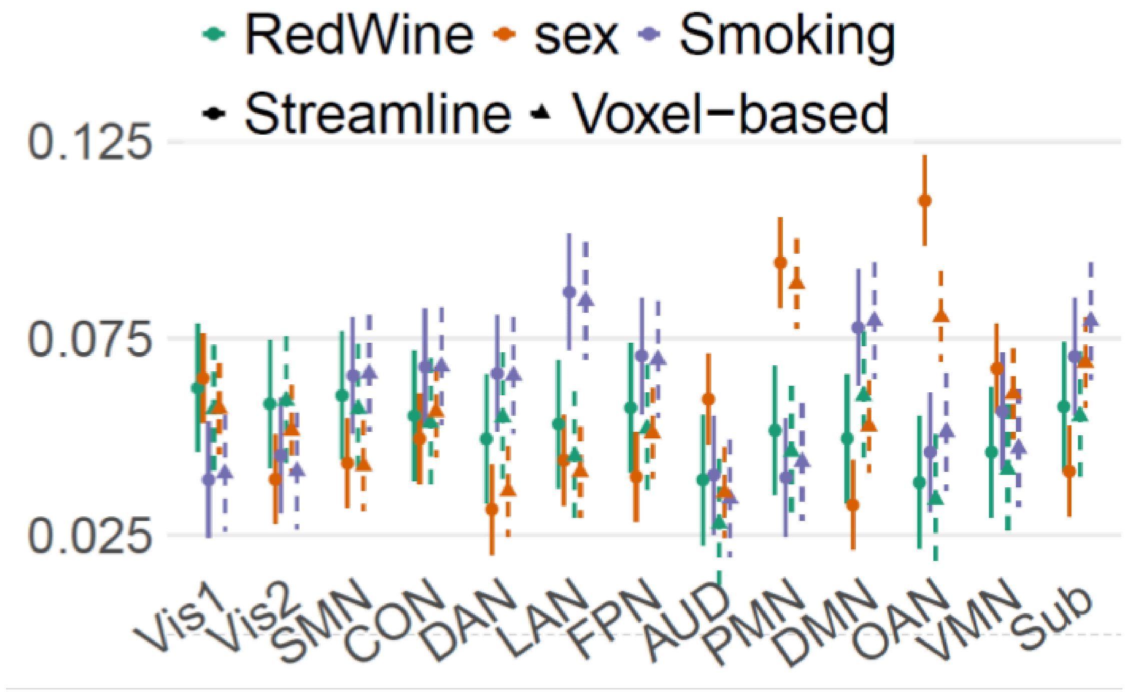


**Supplementary Figure S15. Associations of smoking, red wine consumption and sex with brain-age gap (BAG) predicted from fibers connecting each network, using streamline- and voxel-based features from CBSS, respectively.**

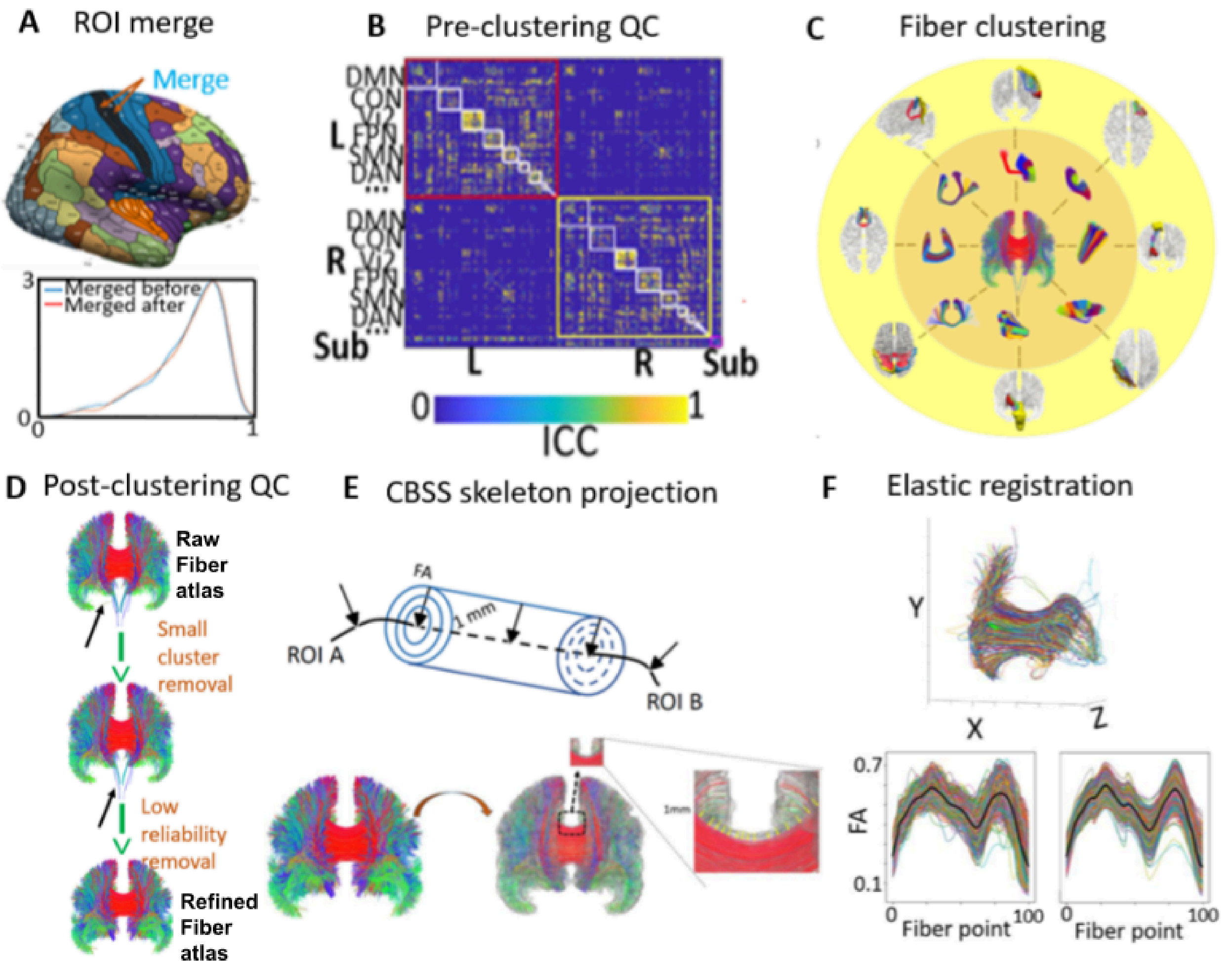


**Supplementary Figure S16**. **Illustrations of the key steps. A.** Merge of ROIs. Upper panel: A merged ROI combining R-3b and R-1 (in black). Lower panel: the distribution density of ICC for fiber counts between all ROI pairs, contrasting between before (blue) and after merging (red). **B.** Pre-clustering QC: ROI pair filtering. ICC matrix to show ICCs of fiber counts between each (merged) ROI pair from 12 networks (white box) in left (red box) and right (yellow) hemisphere and 13 subcortical (pink) ROIs. ROI pairs with fiber count ICC < 0.4 are removed. **C.** Fiber clustering. Fibers between 8 selected ROI pairs are parcelled into clusters. **D.** Post-clustering QC. Clusters with very small membership, or with mean streamline-level ICCs of projected FA values below 0.4 (red), were excluded. **E.** Illustration of 1-mm CBSS fiber-skeleton dilation and fiber projection. **F.** Elastic registration. Elastic registration. Three-dimensional visualization of a representative fiber cluster (top), and FA profiles sampled along its cluster centroid across 1,000 subjects before (bottom left) and after (bottom right) elastic registration. ROI, region of interest; FA, fractional anisotropy; QC, quality control. ICC, intra-class coefficients.

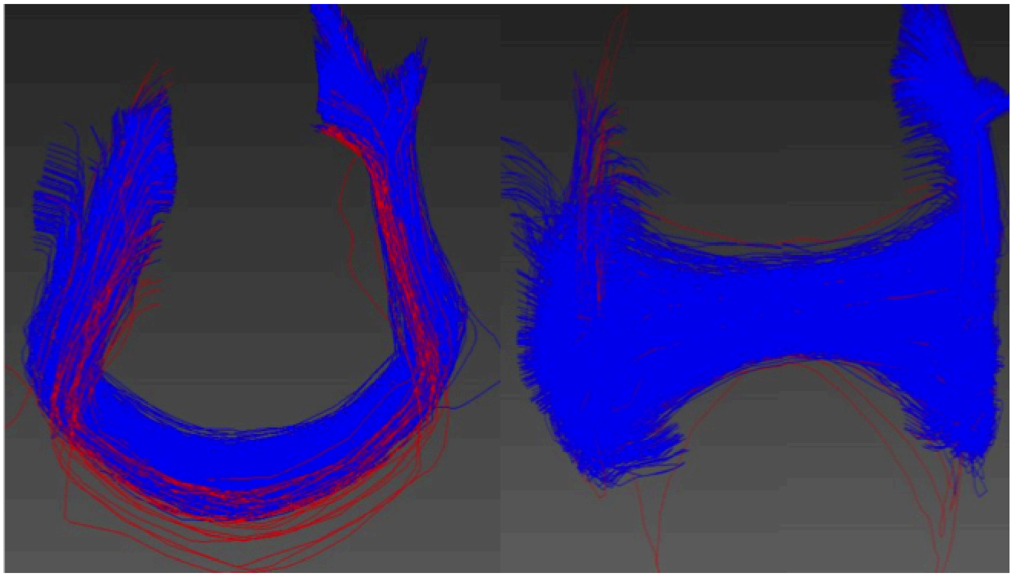

**Supplementary Figure S17**. **Illustrations of the key steps.** Preclustering QC: outlier filtering. Outlier streamlines were identified using QuickBundles in red. QC, quality control.
